# Evasion of APOBEC1-mediated Intrinsic Immunity by a Herpesvirus Uracil DNA Glycosylase Is a Determinant of Viral Encephalitis

**DOI:** 10.1101/2023.06.22.546204

**Authors:** Akihisa Kato, Hayato Harima, Yuji Tsunekawa, Manabu Igarashi, Kouichi Kitamura, Kousho Wakae, Hiroko Kozuka-Hata, Masaaki Oyama, Mizuki Watanabe, Kousuke Takeshima, Yuhei Maruzuru, Naoto Koyanagi, Takashi Okada, Masamichi Muramatsu, Yasushi Kawaguchi

**Author notes:** Address correspondence to: Dr. Yasushi Kawaguchi Division of Molecular Virology Department of Microbiology and Immunology The Institute of Medical Science The University of Tokyo 4-6-1 Shirokanedai, Minato-ku, Tokyo 108-8639, Japan. These authors contributed equally to this work (Akihisa Kato, Hayato Harima).

## Abstract

Herpes simplex virus 1 (HSV-1) is the most common cause of viral encephalitis, which can be lethal or result in severe neurological defects, even when treated with antiviral therapy. We demonstrated that activation of HSV-1 uracil-DNA glycosylase (vUNG) by phosphorylation, essential for its enzymatic activity, counteracted APOBEC1 to promote viral replication and encephalitis in the central nervous system (CNS) of mice. The activation of vUNG protected HSV-1 genomes from APOBEC1-mediated DNA editing, allowing efficient viral replication to occur. The presence of APOBEC1 markedly improved lethal encephalitis in mice infected with an HSV-1 mutant carrying a mutation in the phosphorylation site and an UNG inhibitor protected wild-type HSV-1-infected mice from lethal encephalitis. These findings re-define vUNG as an important factor that allows evasion from intrinsic anti-viral immunity mediated by APOBEC1 in the CNS, and suggest a new therapeutic approach for the treatment of fetal and critical HSV-1 encephalitis.

---

HSV-1, the most prevalent human virus worldwide, causes various mucocutaneous and skin diseases^1^. HSV-1 is also the most common cause of sporadic viral encephalitis, which can be lethal or cause severe neurological defects in survivors, even after antiviral therapy^1, 2^.

The coevolution of viruses and hosts can be characterized as a virus-host arms race whereby hosts have evolved innate, adaptive and intrinsic immunity against viruses to eliminate them, whereas viruses have evolved diverse mechanisms allowing them to survive by evading host responses^3, 4^. Over 400 million years of coevolution, herpesviruses have established a sophisticated balance with their hosts, allowing them to successfully persist over a lifetime and be transmitted to new hosts without causing much damage^5, 6^. Of note, among various diseases caused by HSV-1, encephalitis is unique because it is acute and life-threatening, in contrast to other diseases that manifest recurrently in the periphery^1, 2^. This suggests a delicate balance between virus and host immune response has not been established in the CNS. Therefore, the clarification of interplay between host immune response(s) and viral mechanism(s) to evade host response(s) in the CNS, which promotes viral encephalitis, is critical to understand HSV-1 pathogenesis and develop new therapeutic strategies for HSV-1 encephalitis.

Intrinsic immune responses mediated by endogenous host restriction factors that directly restrict viral replication and assembly are the first line of cellular defense against viral infection^4, 7^. Factors that restrict viral infections include the activation-induced cytidine deaminase and the apolipoprotein B mRNA editing enzyme, catalytic polypeptide-like (AID/APOBEC) family of proteins^8, 9^. Many studies have investigated interplay between host intrinsic immune responses mediated by AID/APOBEC family members and viral evasion mechanisms, especially in retroviruses^8, 9^. However, most studies used cell cultures, and evidence *in vivo* is limited, especially for human pathogenic viruses. Furthermore, there is a lack of information on interplay in the CNS associated with viral encephalitis. Here, we clarified an HSV-1 mechanism that allows evasion from host intrinsic immunity mediated by an AID/APOBEC family protein, which is critical for HSV-1 encephalitis.

We focused on an HSV-1-encoded enzyme, vUNG, which, like cellular UNGs, removes uracil from DNA and initiates base excision repair (BER) to protect DNA from the deleterious consequences of uracil^10, 11^. When examining vUNG regulation in HSV-1-infected cells, vUNG phosphorylation was important for vUNG enzymatic activity in HSV-1-infected cells. Thus, treatment of lysates from wild-type HSV-1(F)-infected HEp-2/ΔhUNGs cells, in which endogenous UNG activity was barely detectable due to the knock-out (KO) of human UNG1 and UNG2, which did not affect cell viability (S-Fig. 1a–e), with phosphatase resulted in the faster electrophoretic mobility of vUNG in denaturing gels and mostly abolished vUNG enzymatic activity in lysates (Fig. 1a–c).

To identify vUNG site(s) whose phosphorylation regulated its enzymic activity, we performed a phosphoproteomic analysis of wild-type HSV-1(F)-infected HEp-2 cells and identified 11 phosphorylation sites (S-Table 1). Of these, serine 302 (Ser-302) was the only residue conserved in UNGs encoded by all human herpesviruses (Fig. 1d). A 100-ns molecular dynamics simulation predicted vUNG Ser-302 phosphorylation promoted flexibility of its DNA intercalation loop, which penetrates into the DNA double helix allowing the insertion of uracil into the binding pocket of the active site^12^, which increases its chance of binding with the vUNG active site (Fig. 1e, S-Fig. 1f, g). Therefore, we focused on Ser-302 phosphorylation in vUNG. vUNG activity in HEp-2/ΔhUNGs cells infected with the recombinant virus vUNG- S302A, in which vUNG Ser-302 was substituted with alanine (S302A) (S-Fig. 2a), but not recombinant virus vUNG-S53A, in which another phosphorylation site vUNG Ser-53 (S-Table 1) was substituted with alanine (S53A) (S-Fig. 2a), was significantly reduced compared with cells infected with wild-type HSV-1(F) or a repaired virus vUNG-SA-repair (Fig. 1f, g and S- Fig. 2a). Similar results were obtained with the recombinant virus vUNG-Q177L/D178N encoding an enzyme-dead mutant of vUNG, in which two amino acids in the activation loop of the enzyme were mutated (S-Fig. 2b)^13^ and its repaired virus vUNG-QL/DN-repair (Fig. 1h, i, and S-Fig. 2a). vUNG activity in HEp-2/ΔhUNGs cells infected with vUNG-S302A was similar to that with vUNG-Q177L/D178N (Fig. 1h, i) indicating Ser-302 phosphorylation in vUNG was essential for vUNG enzymatic activity in HSV-1-infected cells.

In HEp-2 cells infected with wild-type HSV-1(F) or recombinant viruses encoding Flag-tagged wild-type vUNG (vUNG-SA-repair or vUNG-QL/DN-repair) or each of its mutants (S-Fig. 2a), Flag-tagged wild-type vUNGs and Flagg-tagged vUNG-S53A localized diffusely throughout the nucleus and co-localized with the HSV-1 processivity subunit of viral DNA polymerase (vPOL), encoded by the UL42 gene that acts in the BER pathway together with vUNG and vPOL (S-Fig. 3a, c)^11^. In contrast, Flag-tagged vUNG-Q177L/D178N was mislocalized to discrete nuclear domains, termed virus induced chaperon enriched (VICE) domains, that contain Hsc70 and other cellular proteins involved in the proteostatic machinery^14^, and did not co-localize with the vPOL processing factor (S-Fig. 3). Thus, vUNG enzymatic activity is required for proper localization and association with the vPOL processing factor in HSV-1-infected cells. Notably, Flag-tagged vUNG-S302A had a similar localization to Flag-tagged vUNG-Q177L/D178N in HSV-1-infected cells (S-Fig. 3), confirming vUNG phosphorylation is essential for its enzymatic activity in infected cells.

As previously reported with vUNG-null mutant viruses in hamster kidney BHK C13 and mouse fibroblast-like NIH 3T3 cells^10, 15^, growth of vUNG-S302A, vUNG-Q177L/D178N or vUNG-S53A was similar to wild-type HSV-1(F), vUNG-SA-repair or vUNG-QL/DN-repair in Vero cells at multiplicity of infections (MOIs) of 10 or 0.01 (S-Fig. 4a–f). Vero cells infected with vUNG-S302A, vUNG-Q177L/D178N or vUNG-S53A accumulated vUNG at levels similar to cells infected with wild-type HSV-1(F), vUNG-SA-repair or vUNG-QL/DN-repair (S-Fig. 4g, h). Similar results were seen with recombinant viruses encoding Flag-tagged vUNGs (S-Fig. 4i–k), indicating vUNG enzymatic activity and vUNG Ser-302 phosphorylation have no effect on viral replication or vUNG accumulation in cell cultures. In addition, progeny virus yields in HEp-2/ΔhUNGs cells infected with vUNG-S302A or vUNG- Q177L/D178N for 24 h at an MOI of 10 or 48 h at an MOI of 0.01 were similar to infection with wild-type HSV-1(F) (S-Fig. 5), indicating endogenous UNG activity is not required for HSV-1 replication in the absence of vUNG activity in cell cultures, as previously reported for varicella-zoster virus^16^.

To clarify the effects of vUNG activation by Ser-302 phosphorylation in HSV-1 infection *in vivo*, ICR mice were infected ocularly with vUNG-S302A, vUNG-SA-repair, vUNG-Q177L/D178N or vUNG-QL/DN-repair. In this murine model, the capacity to invade the CNS from the peripheral site and to damage the CNS due to viral replication can be studied and the subsequent mortality results from viral encephalitis^17–20^. Survival of mice infected with vUNG-S302A was significantly greater than those infected with vUNG-SA-repair (Fig. 2a). Virus titers in brains of mice infected with vUNG-S302A 1–5 days post-infection were similar to infection with vUNG-SA-repair (Fig. 2b). In contrast, virus titers in brains of mice infected with vUNG-S302A 7 days post-infection were significantly lower than infection with vUNG-SA-repair (Fig. 2b). Similar results were obtained with vUNG-Q177L/D178N and vUNG-QL/DN-repair (Fig. 2c, d), suggesting vUNG Ser-302 phosphorylation is critical for its enzymatic activity in cell culture and *in vivo*. Thus, vUNG activation by Ser-302 phosphorylation has no effect on viral CNS invasiveness or viral replication 5 days post-infection, but is required for efficient viral replication in the CNS 7 days post-infection and viral mortality in mice. Replication of vUNG-S302A and vUNG-Q177L/D178N in brains was reminiscent of an HSV-1 mutant in which UL13, a viral evasion factor for CD8^+^ T cells, was mutated^17^. These suggest vUNG activation by Ser-302 phosphorylation promotes viral replication and pathogenicity by evading host immunity in the CNS.

Enzymatic cytosine deamination, an intrinsic antiviral immune response, by AID/APOBEC family proteins promotes genomic uracil, which is removed by UNG (Fig. 2e)^21^. Therefore, AID/APOBEC family protein(s) induced by HSV-1 infection in mouse brains might restrict viral replication by editing viral DNA genomes, and vUNG might counteract this by initiating the BER pathway. Thus, we examined the effects of HSV-1 infection on mRNA expressions of murine endogenous AID/APOBEC family proteins including AID, APOBEC1, APOBEC2 and APOBEC3 in the brains of ICR mice following ocular infection. mRNA levels of APOBEC1 and APOBEC3, but not AID and APOBEC2, were significantly elevated in brains after vUNG-S302A or vUNG-SA-repair infection (Fig. 2f–i).

Next, we examined the effects of APOBEC1-KO or APOBEC3-KO on vUNG- S302A or vUNG-SA-repair pathogenicity and replication in C57BL/6 mouse brains following intracranial infection. In this murine model, the effects of HSV-1 infection on the CNS can be directly studied and the subsequent mortality results from viral encephalitis^18, 22–24^. As for ICR mice ocularly infected with each recombinant virus, mortality rates and virus titers in wild-type C57BL/6 mouse brains infected with vUNG-S302A or vUNG-Q177L/D178N were significantly lower than for vUNG-SA-repair or vUNG-QL/DN-repair, respectively (S-Fig. 6a–d). The survival rates and virus titers of vUNG-S302A-infected mice were similar to vUNG-Q177L/D178N-infected mice (S-Fig. 6e, f), confirming vUNG Ser-302 phosphorylation was critical for its enzymatic activity *in vivo*.

The survival of APOBEC1-KO mice infected with vUNG-S302A was significantly decreased compared with infected wild-type mice, and similar to wild-type and APOBEC1- KO mice infected with vUNG-SA-repair (Fig. 2j). In contrast, the survival of APOBEC3-KO mice infected with vUNG-S302A was comparable with wild-type mice infected with vUNG- S302A (Fig. 2k). In agreement with the viral pathogenicity of these mice, virus titers in brains of APOBEC1-KO mice infected with vUNG-S302A were significantly increased compared with infected wild-type mice, similar to wild-type and APOBEC1-KO mice infected with vUNG-SA-repair (Fig. 2l). Similar effects of APOBEC1 on vUNG-S302A infection were also observed with vUNG-Q177L/D178N (S-Fig. 6g, h). Thus, APOBEC1 is required for defects in viral replication and CNS pathogenicity caused by the S302A or enzyme-dead mutation in vUNG, suggesting vUNG activation by Ser-302 phosphorylation promotes viral replication and encephalitis by inhibiting APOBEC1-dependent host response(s) against HSV-1 infection. To clarify how vUNG activation by Ser-302 phosphorylation allows evasion from APOBEC1-mediated restriction against HSV-1 infection, HEp-2/ΔhUNGs cells were transfected with an HSV-1 infectious genome clone of vUNG-S302A or vUNG-SA-repair combined with a plasmid expressing human APOBEC1 (hAPOBEC1) fused to EGFP and a self-cleavage site P2A (EGFP-P2A-hAPOBEC1) or its empty plasmid, lysed and subjected to immunoblotting, titration of progeny virus yields, and differential DNA denaturation polymerase chain reaction (3D-PCR), to analyze cytidine deaminase-mediated DNA editing^25, 26^. Transfection of each HSV-1 infectious genome into cells resulted in global viral gene expression and progeny virus production^27^. In 3D-PCR (Fig. 3a), HSV-1 DNA was recovered only at a denaturing temperature of 93.8℃ from lysates of cells expressing vUNG-S302A or hAPOBEC1 and wild-type vUNG (vUNG-SA-repair). In contrast, HSV-1 DNA was recovered at a lower denaturing temperature of 88.9℃ as well as 93.8℃ from lysates of cells expressing hAPOBEC1 and vUNG-S302A. HSV-1 DNA recovery at the lower denaturing temperature indicated a hyper-mutation was induced into the viral genome^25, 26^. Sequence analyses of fragments amplified by PCR verified at 88.9℃ showed they accumulated extensive C-to-T mutations and converted glutamine codon CAG to a TAG stop at two positions in the target HSV-1 Us3 gene (Fig. 3b and S-Table 2). In contrast, fragments amplified at 93.8°C had few mutations (S-Fig. 7a–c and S-Table 2). Progeny virus yields and expression levels of vUNG and another viral protein glycoprotein B (gB) from lysates of cells expressing hAPOBEC1 and vUNG-S302A were significantly lower in cells expressing vUNG-S302A or hAPOBEC1 and wild-type vUNG (Fig. 3c–g). Thus, reduced progeny virus yields were linked to hyper- mutation in the viral genome. We verified that upon ectopic expression, human and mouse APOBEC1 were localized in the nucleus of HSV-1-infected HEp-2 cells (S-Fig. 7d, e), where viral DNA replication occurs^1^, and that HSV-1 infection did not induce detectable levels of endogenous hAPOBEC1 by immunoblotting of HEp-2 cells (data not shown). Thus, hAPOBEC1 impairs viral genome integrity by inducing hyper-mutation in the absence of vUNG enzymatic activity, thereby reducing mutant virus (vUNG-S302A) replication. This suggests vUNG activated by Ser-302 phosphorylation counteracts impaired viral genome integrity and viral replication caused by APOBEC1.

To address the potential of vUNG as a therapeutic target for HSV-1 encephalitis, wild-type or APOBEC1-KO mice pretreated with an adeno-associated virus (AAV) vector expressing UGI (AAV-UGI), an inhibitor of many UNGs including vUNG (Fig. 4a)^28^, or a control AAV vector expressing a fluorescence protein ZsGreen AAV-ZsGreen, were intracranially infected with vUNG-SA-repair and their survival was analyzed. We verified ZsGreen fluorescence was detected throughout brains (S-Fig. 7f), confirming the high efficiency of AAV transduction in the brains of mice. Treatment with AAV-UGI, but not AAV-ZsGreen, significantly improved the survival rate of wild-type mice but not mAPOBEC1-KO mice (Fig. 4b). Thus, UGI effectively protected mice from lethal encephalitis dependent on APOBEC1.

Virally-encoded UNGs have been postulated to maintain viral genome integrity by removing genomic uracil during viral replication^11^. Uracil in DNA occurs by spontaneous hydrolytic cytosine deamination, enzymatic cytosine deamination with AID/APOBEC family proteins, and thymine replacing misincorporation (Fig. 2e)^21^. Currently, interplay between viral UNGs and AID/APOBEC family proteins during viral infection is unknown. Therefore, the route targeted by viral UNGs is unclear. We showed APOBEC1 was induced by HSV-1 infection in the CNS of mice and the activation of vUNG by Ser-302 phosphorylation, which is essential for its enzymatic activity, counteracted APOBEC1-mediated intrinsic immunity, thereby promoting viral replication and encephalitis in the CNS. vUNG activation by phosphorylation had no effect on viral replication and pathogenicity in APOBEC1 KO mice (Fig. 2j, l), suggesting vUNG mainly acts against APOBEC1 but not spontaneous hydrolytic cytosine deamination or thymine replacing uracil misincorporation *in vivo*. Thus, we re-defined HSV-1-encoded UNG as a significant viral evasion factor against APOBEC1 and possibly other AID/APOBEC family proteins. This is the first study to report interplay between intrinsic immunity and viral evasion mechanisms in the CNS critical for viral encephalitis, and that APOBEC1 is an intrinsic anti-viral immune factor *in vivo*. These *in vivo* observations are of importance, based on the effects of AID/APOBEC family proteins on viral infection in cell cultures, especially by overexpression of restriction factors, which often contradict *in vivo* results^29–33^. Notably, the Allen human brain atlas database^34^ indicates that APOBEC1 is expressed in the human brain (S-Fig. 8).

Protection of mice infected with the HSV-1 mutant carrying the S302A mutation against lethal encephalitis was associated with APOBEC1 (Fig. 2j) and the effective protection of HSV-1-infected mice against lethal encephalitis by the UNG inhibitor UGI (Fig. 4b) demonstrated the inhibitory effect of APOBEC1 on HSV-1 infection in the CNS is effective in the absence of vUNG. Thus, evasion from APOBEC1 by vUNG is critical for HSV-1 infection in the CNS, which suggests a new therapeutic approach for the treatment of fetal and critical HSV-1 encephalitis—restoration of intrinsic anti-viral immunity by inhibiting the viral evasion factor. These therapeutic possibilities suggest the development of drugs that inhibit vUNG or a kinase responsible for the phosphorylation of vUNG at Ser-302. The development of drugs that specifically inhibit viral UNGs but not human UNGs has been continuously attempted^35, 36^.

Currently, whether UNGs encoded by poxviruses and other herpesviruses, like HSV- 1 UNG, counteract AID/APOBEC family members during infection remains unclear. Epstein-Barr virus (EBV), another human herpesvirus, has evolved an evasion factor BORF2, which directly binds, inhibits and relocalizes APOBEC3B in cell cultures^26^. Furthermore, mutations indicative of human APOBEC3 editing were recently detected in a specific clade of monkey pox viruses isolated from humans^37^. These observations suggest poxviruses and other herpesviruses are potential substrates for AID/APOBEC family members and viral UNGs may counteract their restrictions. In contrast, APBEC3G overexpression had no effect on infection of another pox virus, vaccinia virus, in cell cultures^38^. Thus, UNG encoded by this virus might antagonize the effect of APOBEC3G on viral infection as observed in this study with APOBEC1 (Fig. 3).

## METHODS

### Cells and viruses

Simian kidney epithelial Vero, human carcinoma HEp-2 and rabbit skin cells as well as HSV-1 wild-type strain HSV-1(F) were described previously^27, 39, 40^. Plat*-* GP cells from a 293T-derived murine leukemia virus-based packaging cell line were described previously^41^. HEK293EB cells stably expressing the E1 gene region and BCL-XL gene^42^ were maintained in DMEM containing 8% FCS.

**Figure 1.**
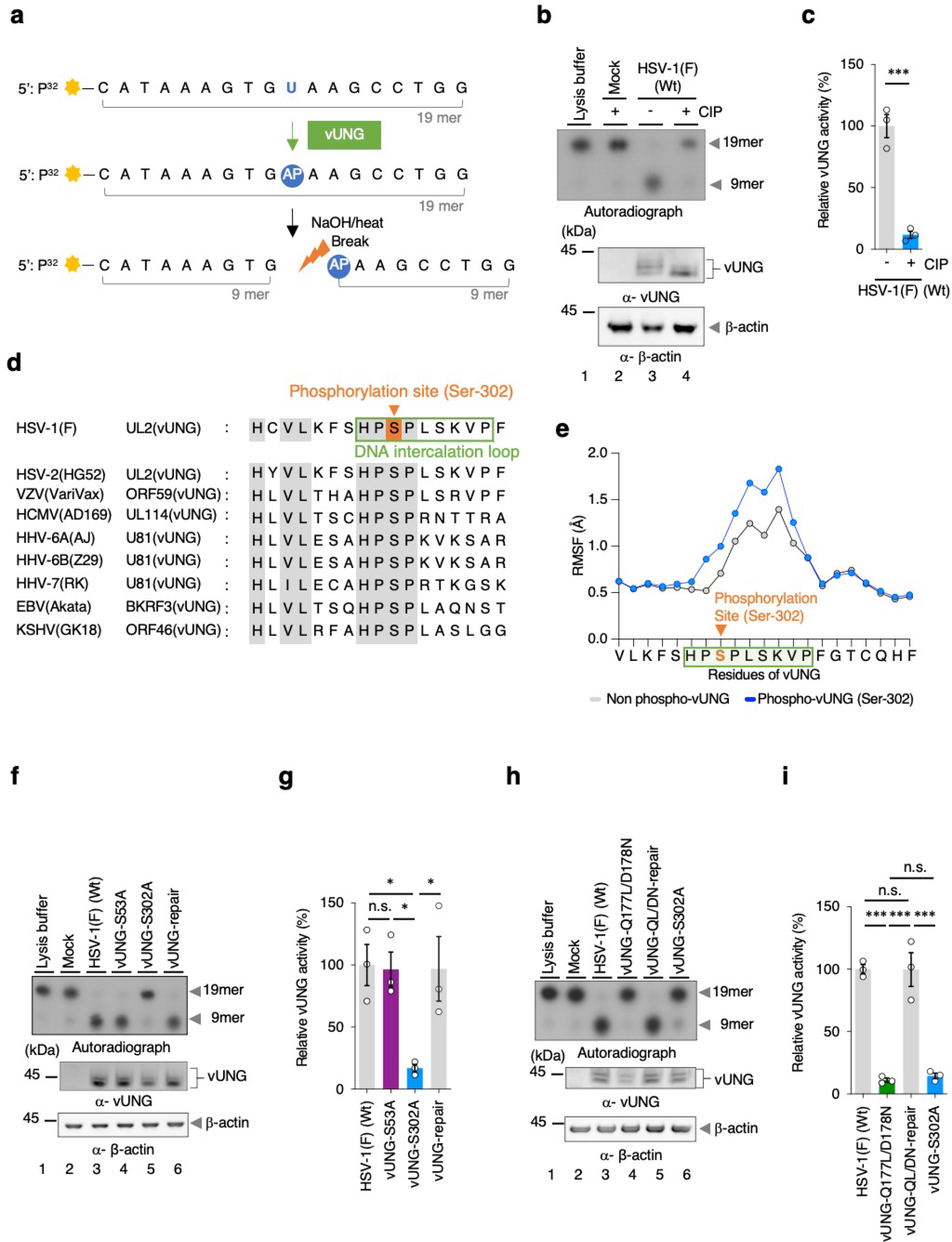
**Phosphorylation of vUNG Ser-302 is essential to activate vUNG enzymatic activity. a**. Flow chart for the vUNG assay. P^32^-labeled 19-mer oligonucleotides containing an uracil residue at position 10 were used as substrates. Following the vUNG reaction, reaction products were treated with 0.4 N NaOH/heat to cleave AP sites generated by vUNG. Final products were displayed by 20% polyacrylamide/7M urea gels. **b**. Uracil excision activity of lysis buffer (lane 1), lysates from mock-infected HEp-2/ΔhUNGs cells (lane 2) and untreated cells infected with wild-type HSV-1(F) for 24 h at an MOI of 10 (lane 3), or treated with CIP (lane 4) (upper panel). The cell lysates prepared for the vUNG assays were analyzed by immunoblotting with antibodies to vUNG (middle panel) or Δ-actin (lower panel). **c**. The amounts of cleaved oligonucleotides in the experiment in (**b**) were quantitated and normalized to those of vUNG proteins. Each value is the mean ± SEM of three independent experiments and is expressed relative to that in untreated HSV-1(F)-infected cells (lane 3 in (**b**)), which was normalized to 100%. The statistical significance determined by an unpaired two-tailed Student’s *t*-test is indicated. ***; *p* < 0.001. **d**. Amino acid sequence alignment of UL2 (vUNG) homologs from HSV-1(F), HSV-2(HG52), VZV(VariVax), HCMV(AD169), HHV-6A(AJ), HHV-6B(Z29), HHV-7(RK), EBV(Akata) and KSHV(GK18). Residues conserved in the sequences and phosphorylation sites are shaded gray and orange, respectively. Amino acid residues of the DNA intercalation loop of HSV-1 vUNG are shown in a green square. **e**. The root-mean-square-fluctuation (RMSF) analysis of non-phosphorylated and phosphorylated vUNG during the MD simulation period of 100 ns in the experiment in S-Fig. 1g. Amino acid residues of DNA intercalation loop are shown in a green square. **f** to **i**. HEp-2/ΔhUNGs mock-infected (**f**, **h**) or infected with wild-type HSV-1(F) (**f**, **h**), vUNG-S53A (**f**), vUNG-S302A (**f**, **h**), vUNG-SA-repair (**f)**, vUNG-Q177L/D178N (**h**), or vUNG-QL/DN-repair (**h**) were analyzed by vUNG assay (upper panel) or immunoblotting with antibodies to vUNG (middle panel) or Δ-actin (lower panel). The amounts of cleaved oligonucleotides in the experiments in (**f**) and (**h**) were quantitated and normalized to those of vUNG proteins. Each value is the mean ± SEM of three independent experiments and is expressed relative to that in HSV-1(F)-infected cells, which was normalized to 100%. The statistical significance determined by one-way ANOVA followed by Tukey’s test is indicated. *; *p* < 0.05, ***; *p* < 0.001. n.s. not significant.

**Figure 2.**
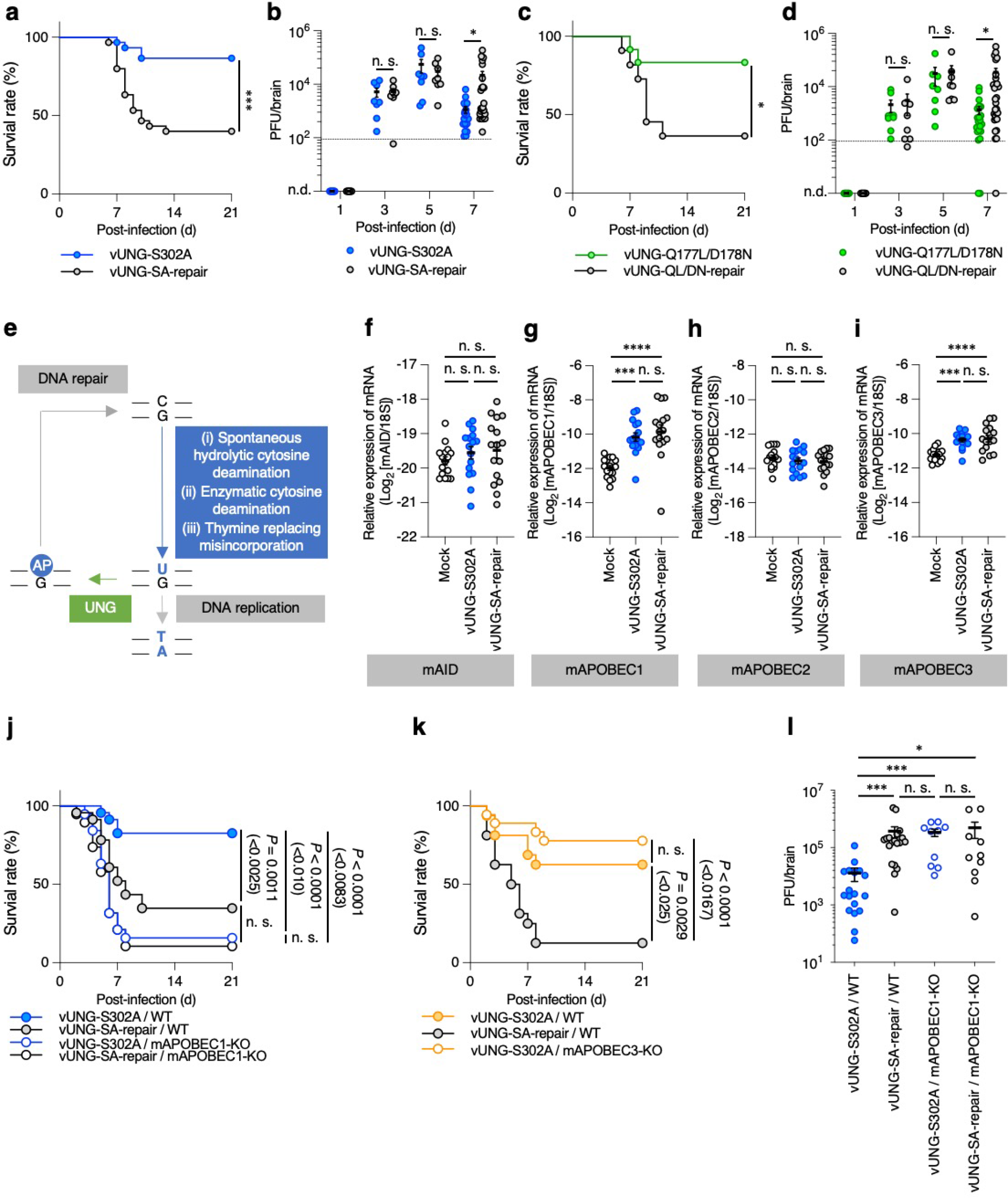
**Counteraction of APOBEC1 by vUNG is critical for viral replication and pathogenicity in the CNS of mice. a**, **c**, Four-week-old female ICR mice were infected ocularly with 3 × 10^6^ PFU/eye of vUNG-S302A (*n* = 30) (**a**), vUNG-SA-repair (*n* = 30) (**a**), vUNG-Q177L/D178N (*n* = 12) (**c**) or vUNG-QL/DN-repair (*n* = 11) (**c**) and monitored daily for survival for 21 days. The statistical significance determined by Log-rank test is indicated. *; *p* < 0.05, ***; *p* < 0.001. **b**, **d**, Viral titers in the brains of mice infected ocularly with 3 × 10^6^ PFU/eye of vUNG-S302A (**b**), vUNG-SA-repair (**b**), vUNG-Q177L/D178N (**d**) or vUNG- QL/DN-repair (**d**) at 1 (*n* = 8), 3 *(n* = 8), 5 (*n* = 8), and 7 days (vUNG-S302A, *n* = 22; vUNG- SA-repair, *n* = 27; vUNG-Q177L/D178N, *n* = 23; vUNG-QL/DN-repair, *n* = 29) post-infection were assayed. Dashed line indicates the limit of detection. n.d. not detected. The statistical significance determined by an unpaired two-tailed Student’s *t-*test (at 3 days in (**b**) and (**d**); 5 and 7 days in (**d**)) and Welch’s *t*-test (at 5 and 7 days in (**b**)) is indicated. *; *p* < 0.05, n.s. not significant. **e**, Working flow diagram of the development and repair of genomic uracil. **f** to **i**, The amounts of mAID, mAPOBEC1, mAPOBEC2 and mAPOBEC3 mRNA in the brains of 4-week-old female ICR mice mock-infected or infected ocularly with 3 × 10^6^ PFU/eye of vUNG-S302A or vUNG-SA-repair (*n* = 16). Each value is the mean ± SEM for each group. The statistical significance determined by one-way ANOVA followed by Tukey’s test is indicated. ***; *p* < 0.001, ****; *p* < 0.0001, n.s., not significant. **j**, **k**, Three-to-six-week-old C57BL/6 WT (*n* = 23) (**j**), APOBEC1-KO (n = 19) (**k**), WT (*n* = 16) (**j**) or APOBEC3-KO (*n* = 18) (**k**) mice were infected intracranially with 1×10^3^ PFU/head of vUNG-S302A or vUNG- SA-repair, and monitored daily for survival for 21 days. Statistical analysis was performed by Log-rank test, and for four (**j**) or three (**k**) comparison analyses, *P-*values <0.0083 (0.05/6), <0.01 (0.05/5), <0.0125 (0.05/4), <0.0167 (0.05/3), <0.025 (0.05/2), or <0.05 (0.05/1) were sequentially considered significant after Holm’s sequentially rejective Bonferroni multiple- comparison adjustment. **l**, Three-to-six-week-old C57BL/6 WT or APOBEC1-KO mice were infected intracranially with 1×10^3^ PFU/head of vUNG-S302A or vUNG-SA-repair. Viral titers in the brains of infected mice at 5 days post-infection were assayed. Each data point is the virus titer in the brain of one infected mouse: WT mice were infected with vUNG-S302A (*n* = 18) or vUNG-SA-repair (*n* = 19), and APOBEC1-KO mice were infected with vUNG-S302A (*n* = 10) or vUNG-SA-repair (*n* = 10). The statistical significance determined by one-way ANOVA followed by Tukey’s test is indicated. *; *p* < 0.05, ***; *p* < 0.001, n.s., not significant.

**Figure 3.**
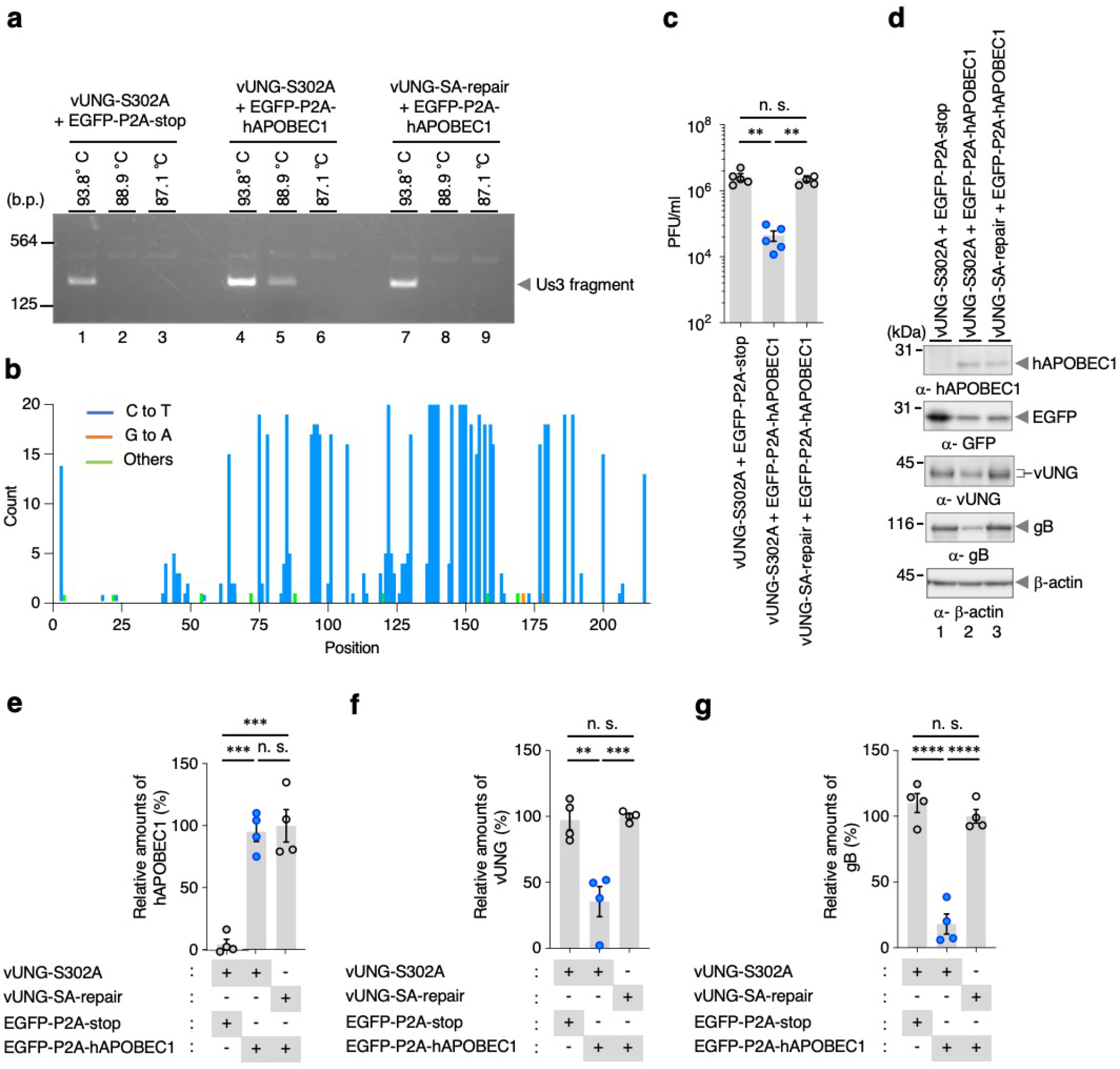
**hAPOBEC1 inhibits viral replication by editing the HSV-1 genome. a**, HEp- 2/ΔhUNGs cells were co-transfected with an HSV-1 infectious genome clone of vUNG-S302A (lanes 1 to 6) or that of vUNG-SA-repair (lanes 7 to 9) along with pcDNA-EGFP-P2A-stop (lanes 1 to 3) or pcDNA-EGFP-P2A-hAPOBEC1 (lanes 4 to 9). At 6 days post-transfection, total DNA purified from each cell was subjected to Us3 3D-PCR assays. Arrows indicate the position of the 3D-PCR products (257 bp). Digital images are representative of three independent experiments. **b**, In the experiment in (**a**), Us3 DNA amplified at a denaturation temperature of 88.9°C (lane 5) was excised, cloned, and sequenced. Results are summarized: C-to-T and G-to-A hypermutations are indicated as blue and orange vertical lines, respectively. All other base substitutions are shown as green vertical lines. **c**, HEp-2/ΔhUNGs cells were co- transfected with an HSV-1 infectious genome clone of vUNG-S302A or that of vUNG-SA- repair along with pcDNA-EGFP-P2A-stop or pcDNA-EGFP-P2A-hAPOBEC1. At 6 days post-transfection, viral titers of each cell were assayed. Each value is the mean ± SEM for five independent experiments. The statistical significance determined by one-way ANOVA followed by Tukey’s test is indicated. **; *p* < 0.01, n.s., not significant. **d**, HEp-2/ΔhUNGs cells co-transfected with an HSV-1 infectious genome clone of vUNG-S302A (lanes 1 and 2) or that of vUNG-SA-repair (lane 3) along with pcDNA-EGFP-P2A-stop (lane 1) or pcDNA-EGFP- P2A-hAPOBEC1 (lanes 2 and 3) for 6 days were analyzed by immunoblotting with antibodies to APOBEC1, GFP, vUNG, gB, or Δ-actin. Digital images are representative of four independent experiments. **e**, **f**, **g**, The amounts of hAPOBEC1 (**e**), vUNG (**f**) and gB (**g**) proteins in the experiment in (**d**) were quantitated and normalized to those of Δ-actin proteins. Each value is the mean ± SEM of four independent experiments and is expressed relative to that in cells co-transfected with the HSV-1 infectious genome clone of vUNG-SA-repair and pcDNA-EGFP-P2A-hAPOBEC1, which was normalized to 100%. The statistical significance determined by one-way ANOVA followed by Tukey’s test is indicated. **; *p* < 0.01, ***; *p* < 0.001, ****; *p* < 0.0001, n.s., not significant.

**Figure 4.**
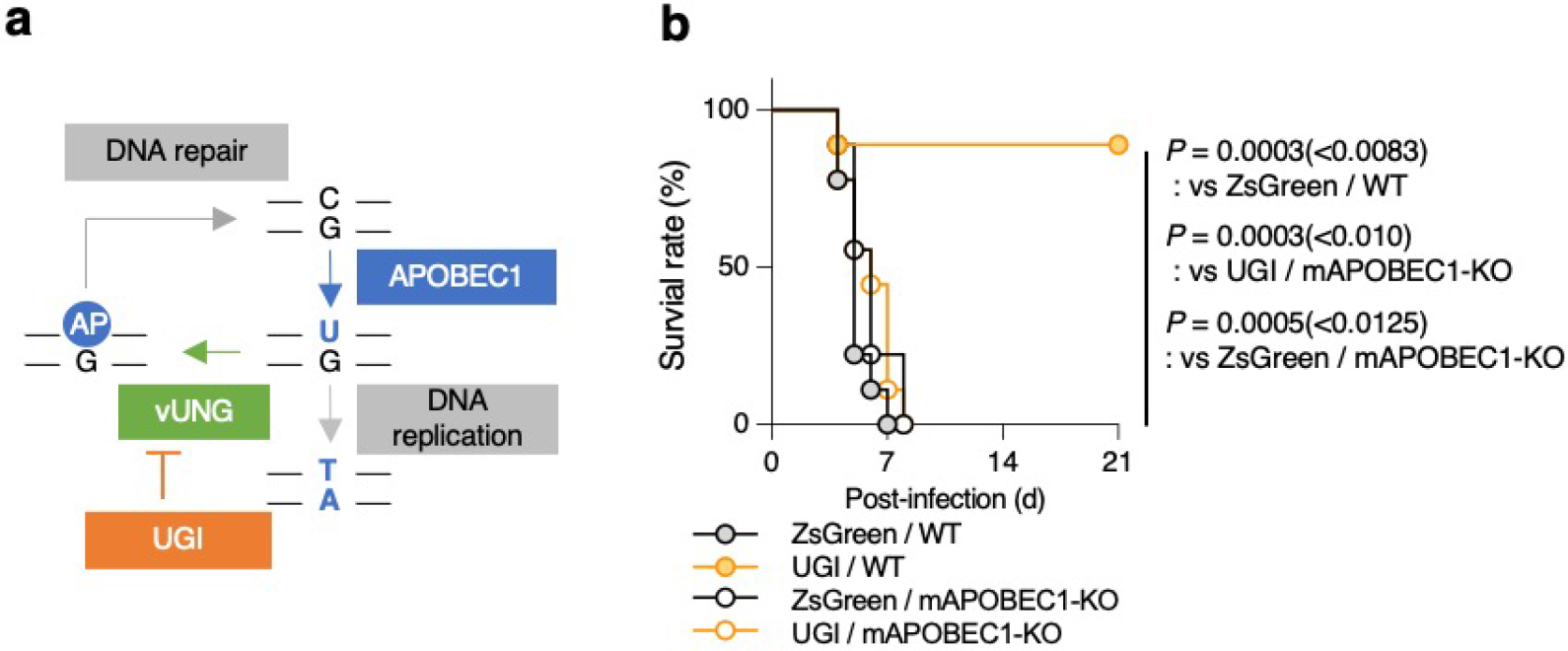
**UGI protects mice from lethal encephalitis in a manner dependent on APOBEC1. a**, Working flow diagram of the development and repair of genomic uracil, and UGI action point. **b**, Three-week-old C57BL/6 WT or APOBEC1-KO mice were pretreated intracranially with 5×10^10^ vg/head of AAV-UGI or AAV-ZsGreen, respectively. At 14 days after pretreatment with AAV-vectors, the mice were infected intracranially with 2×10^4^ PFU/head of vUNG-SA-repair, and monitored daily for survival for 21 days (*n* = 9). Statistical analysis was performed by Log-rank test, and for four comparison analyses, *P-*values < 0.0083 (0.05/6), <0.01 (0.05/5), <0.0125 (0.05/4), <0.0167 (0.05/3), <0.025 (0.05/2), or <0.05 (0.05/1) were sequentially considered significant after Holm’s sequentially rejective Bonferroni multiple- comparison adjustment.

### Mice

Four-week-old female ICR mice were purchased from Charles River for ocular infection. C57BL/6 APOBEC1-KO and APOBEC3-KO mice were provided by K. Yusa (Wellcome Trust Sanger Institute) and E. Jefferies (MRC Laboratory of Molecular Biology), respectively, and were maintained at the Institute of Medical Science, The University of Tokyo. Three-to-six-week-old C57BL/6 mice (JAX: 000664) purchased from Charles River were used as WT controls. APOBEC1-KO, APOBEC3-KO and WT control mice were bred and maintained under conventional conditions. All animal experiments were carried out in accordance with the Guidelines for the Proper Conduct of Animal Experiments of the Science Council of Japan. The protocol was approved by the Institutional Animal Care and Use Committee, Institute of Medical Science, the University of Tokyo (IACUC protocol approvals, PM27-113, PM27-73, PA17-67, PH2-12, A21-55, and A22-27).

### Plasmids

To construct pX330-hUNGs, sense and antisense oligonucleotides were designed for insertion into the *Bbs*I site in a pX330 bicistronic expression vector, which expresses Cas9 and a synthetic single-guide RNA (Addgene), as follows: 5′- CACCGCTGGGCGGAGGCGGAAC-3′ and 5′-AAACGTTCCGCCTCCGCCCAGC-3′. The DNA oligonucleotides were annealed and incorporated into the pX330 vector linearized with the *Bbs*I restriction enzyme. pcDNA-EGFP-P2A-stop was constructed by amplifying the EGFP and P2A sequences from pEGFP-C1 (Clontech) using the primers shown in S-Table 3 and cloning it into the *Bam*HI and *Eco*RI sites of pcDNA3.1(+) (Thermo Fisher Scientific). p3flag- hA1 was constructed by cloning the *Xho*I-*Apa*I fragments of pCMV-hA1^43^ into the *Xho*I and *Apa*I sites of pCMV-3Tag-1C (Agilent). pcDNA-EGFP-P2A-hAPOBEC1 was constructed by amplifying the entire coding sequence of human APOBEC1 (hAPOBEC1) from p3flag-hA1 by PCR using the primers listed in S-Table 3, and cloning it into the *Eco*RI and *Eco*RV sites of pcDNA-EGFP-P2A in frame with the tag sequence by using an In-Fusion® HD Cloning Kit according to the manufacturer’s instructions (Takara). pcDNA-hAPOBEC1 was constructed by cloning the *Xho*I-*Hin*dIII fragments of pUC57-APOBEC1, which was synthesized by GenScript, into the *Eco*RI and *Not*I sites of pcDNA3.1(+) (Thermo Fisher Scientific). pEGFP- mA1 was constructed by cloning the *EcoR*I-*Xho*I fragments of pGEM-mA1, generated by cloning the entire coding sequence of mouse APOBEC1 (mAPOBEC1) amplified from mouse cDNA by PCR into pGEM-T-easy (Promega), and cloning it into the *Eco*RI and *Sal*I sites of pEGFP-C2 (Clontech). pRetroX-TRE3G-hAPOBEC1-3xflag and pRetroX-TRE3G- mAPOBEC1-3xflag were constructed by cloning the entire coding sequences of hAPOBEC1 and mAPOBEC1 amplified from pcDNA-hAPOBEC1 and pEGFP-mA1, respectively, by PCR using the primers shown in S-Table 3 into the *Eco*RI and *Bam*HI sites of pRetroX-TRE3G (TaKaRa). pBS-UGI-flag was constructed by amplifying the UGI and flag sequence from UGI- pFERp^44^ by PCR and cloning it into the *Not*I and *Eco*RI sites of pBluescript II KS(+) (Stratagene). pAAV-UGI was constructed by cloning the entire coding sequence of UGI amplified from pBS-UGI-flag by PCR using the primers shown in S-Table 3 into the *Eco*RI and *Bam*HI sites of pAAV-ZsGreen1 (TaKaRa).

### Establishment of HEp-2/ΔhUNGs cells

HEp-2 cells were transfected with pX330- hUNGs using a NEPA21 electroporator (NepaGene), cloned from a single colony, and designated HEp-2/ΔhUNGs cells. To determine the genotypes of each allele from HEp- 2/ΔhUNGs cells, genomic DNA from these cells was amplified by PCR and sequenced directly. The sequencing of PCR products showed mixed patterns of sequences, and therefore, PCR products were cloned into plasmids, and their sequences were determined. We obtained two patterns of sequences (S-Fig. 1a), which represented the common sequences for hUNG1 and hUNG2 of the two alleles of hUNGs from HEp-2/ΔhUNGs cells and did not include a pattern of the wild-type sequence.

### Assay for cell viability

The viabilities of HEp-2 and HEp-2/ΔhUNGs cells were assayed using a cell counting kit-8 (Dojindo) according to the manufacturer’s instructions.

### Construction of recombinant HSV-1s

Recombinant viruses vUNG-S53A or vUNG-S302A, in which the serine at residue Ser-53 or -302 of vUNG were substituted with alanine, respectively (S-Fig. 2a), were generated by a two-step Red-mediated mutagenesis procedure using *E. coli* GS1783 strain containing pYEbac102Cre^27, 45^ as described previously^46, 47^, except using the primers listed in S-Table 4. The glutamine at residue Gln-177 and aspartic acid at residue Asp-178 in the water-activation loop of HSV-1 vUNG are highly conserved in various UNGs (S-Fig. 2b) and have been reported to be critical for the enzymatic activity of Epstein-Barr Virus UNG^13^. A recombinant virus vUNG-Q177L/D178N, in which the enzymatic activity of vUNG was inactivated by replacing both Gln-177 and Asp-178 with a leucine and asparagine, respectively (S-Fig. 2a), was generated by the two-step Red-mediated mutagenesis procedure using *E. coli* GS1783 strain containing pYEbac102Cre^27, 45^ as described previously^46, 47^, except using the primers listed in S-Table 4. The recombinant viruses Flag- vUNG-S53A, Flag-vUNG-S302A and Flag-vUNG-Q177L/D178N, encoding flag-tagged vUNG and carrying the S53A, S302A and Q177L/D178N mutations, respectively (S-Fig. 2a), were generated by the two-step Red-mediated mutagenesis procedure^46, 47^, except using the primers listed in S-Table 4 and *E. coli* GS1783 containing the vUNG-S53A, vUNG-S302A or vUNG-Q177L/D178N genomes. The recombinant viruses vUNG-SA-repair, vUNG-QL/DN- repair, Flag-vUNG-SA-repair or Flag-vUNG-QL/DN-repair, in which the S302A and Q177L/D178N mutations in vUNG or flag-tagged vUNG were repaired (S-Fig. 2a), were generated by the two-step Red-mediated mutagenesis procedure^46, 47^, except using the primers listed in S-Table 4 and *E. coli* GS1783 containing the vUNG-S302A, vUNG-Q177L/D178N, Flag-vUNG-S302A or Flag-vUNG-Q177L/D178N genomes.

### UNG and vUNG assays

UNG and vUNG activities were determined by measuring the alkaline cleavage of uracil-containing oligonucleotides (Fig. 1a). Briefly, HEp-2 and/or HEp-2/ΔhUNGs cells were mock-infected or infected with wild-type HSV-1(F) and/or each recombinant virus at an MOI of 10. Infected cells were harvested at 24 h post-infection and solubilized in lysis buffer (10 mM Tris-HCl [pH 7.4], 1 mM EDTA, 0.5% Triton X-100, 1 mM DTT) containing a protease inhibitor cocktail (Nacalai Tesque). After brief sonication and centrifugation, 20 μl of supernatant was mixed with 80 μl reaction buffer (10 mM Tris-HCl [pH 7.4], 1 mM EDTA, 62.5 mM NaCl) and 0.5 μl of 0.5 pmol/μl 5’-P^32^ labeled uracil- containing oligonucleotides (5’-CATAAAGTG-U-AAAGCCTGG-‘3), which contains a U at position 10. The reaction was allowed to proceed for 30 min at 37°C and then terminated by adding 75 μl of stop buffer (70% formamide, 0.4 M NaOH, 1x TBE) and by heating at 100°C for 15 min. Subsequently, samples were combined with 35 μl of dye buffer (36% glycerol, 30 mM EDTA, 0.05% bromophenol blue, and 0.035% xylene cyanol), and analyzed by electrophoresis in 20% polyacrylamide/7M urea gels in glycerol tolerant gel buffer (5% glycerol, 40% methanol, 10% acetic acid). The gels were dried and subjected to autoradiography. The amounts of cleaved oligonucleotides was quantified with ImageQuant (GE Healthcare). In the vUNG assay, vUNG activity was normalized to the amount of vUNG proteins present.

### Phosphatase treatment

Lysates of HEp-2/ι1hUNGs cells mock infected or infected with wild-type HSV-1(F) at an MOI of 10 for 24 h were treated with alkaline phosphatase (CIP) (New England BioLabs) as described previously^48^.

### Sample preparation for MS

HEp-2 cells were infected at an MOI of 10 with wild-type HSV-1(F), harvested at 36 h post-infection, and suspended in 8 M urea containing PhosSTOP (Roche Diagnostics) and Benzonase (Novagen). The mixture was kept on ice for 1 h and cellular debris was pelleted by centrifugation at 15,000 rpm for 30 min. The cell lysate was reduced with 1 mM ditiothreitol (DTT) for 90 min and then alkylated with 5.5 mM iodoacetamide (IAA) for 30 min. After digestion with lysyl endopeptidase (Lys-C) (1:50 w/w) (Wako) at 37°C for 3 h, the resulting peptide mixtures were diluted with 10 mM Tris-HCl (pH 8.2) to a final urea concentration ˂ 2 M and then digested with modified trypsin (1:50 w/w) (Sequencing Grade, Promega) at 37°C for 3 h. An equal amount of trypsin was then added overnight for digestion. Phosphopeptides were enriched using a Titansphere Phos-TiO Kit (GL Sciences) as described previously^49^, except that the captured peptides were eluted with a 5% ammonium solution or 5% pyrrolidine solution. The enriched phosphopeptide solutions were acidified with 10% TFA, desalted with ZipTip C18 resins (Millipore) and centrifuged in a vacuum concentrator.

### Mass spectrometric analysis, protein identification and determination of phosphorylated sites

Shotgun proteomic analyses of the Titansphere eluates were performed by a linear ion trap-orbitrap mass spectrometer (LTQ-Orbitrap Velos, Thermo Fisher Scientific) coupled to a nanoflow LC system (Dina-2A, KYA Technologies). Samples mixed with stable-isotope-labeled phospho-peptides, FSHP(p)SPLSK [^13^C and ^15^N-labeled lysine] (Sigma), were injected into a 75 µm reversed-phase C18 column at a flow rate of 10 µl/min and eluted with a linear gradient of solvent A (2% acetonitrile and 0.1% formic acid in H2O) to solvent B (40% acetonitrile and 0.1% formic acid in H2O) at 300 nl/min. Peptides were sequentially sprayed from a nanoelectrospray ion source (KYA Technologies) and analyzed by collision-induced dissociation (CID). The analyses were performed in the data-dependent mode, switching automatically between MS and MS/MS acquisition. For the CID analyses, full-scan MS spectra (from m/z 380 to 2000) were acquired in the orbitrap with a resolution of 100,000 at m/z 400 after ion count accumulation to the target value of 1,000,000. The 20 most intense ions at a threshold above 2000 were fragmented in the linear ion trap with a normalized collision energy of 35% for an activation time of 10 ms. The orbitrap analyzer was operated with the “lock mass” option to perform shotgun detection with high accuracy^50^. Protein identification was conducted by analyzing the MS and MS/MS data against the RefSeq human protein database combined with the virus protein sequences based on the complete genome sequence of HSV-1 by Mascot (Matrix Science). Carbamidomethylation of cysteine residues was set as a fixed modification, whereas methionine oxidation, protein N-terminal acetylation, pyro-glutamination for N-terminal glutamine and phosphorylation (Ser, Thr, and Tyr) were set as variable modifications. A maximum of two missed cleavages was allowed in the database search. The tolerance for mass deviation was set to 3 parts per million for peptide masses and 0.8 Da for MS/MS peaks, respectively. For peptide identification, we conducted decoy database searching by Mascot and applied a filter for a false positive rate < 1%. Determination of phosphorylated sites in peptides was performed using Proteome Discoverer ver 1.3 (Thermo Fisher Scientific).

### Molecular dynamic (MD) simulations

The initial structure of vUNG was obtained from the Protein Data Bank (PDB ID:1UDG). The structure of phosphorylated vUNG (Ser- 302) was generated using BIOVIA Discovery Studio 2017 (Dassault systems). MD simulations of the wild-type and phosphorylated forms of vUNG were performed using the AMBER 16 package^51^. The AMBER ff14SB force field for proteins and phosaa10 parameters for phosphoserines (SEP) were used^52^. The protonation states of the ionizable residues were assigned at pH 7.4 using the PDB2PQR web server^53^. Missing hydrogen atoms were added to the LEaP module in AMBER 16. The total charges were neutralized by the addition of chloride counterions. The systems were then solvated with TIP3P water molecules and 0.15 M NaCl. Energy minimizations and MD simulations were performed using the pmemd.cuda program in AMBER 16, with a cutoff radius of 10 Å for the nonbonded interactions. All systems were energy-minimized in two steps: first, water and ions; second, all atoms. For energy minimization, the steepest descent method was used for 500 steps followed by the conjugate gradient method for 1500 steps. After energy minimization, the system was gradually heated from 0 K to 310 K over 300 picoseconds (ps) with harmonic restraints (with a force constant of 1.0 kcal/mol·Å^2^). Two additional rounds of MD simulations (50 ps each at 310 K) were performed with decreasing restraint weight from 0.5 to 0.1 kcal/mol·Å^2^. Next, unrestrained production runs of 100 nanoseconds (ns) for vUNG were performed, and the production trajectories were collected every 10 ps. All MD simulations were performed using the NPT ensemble and Berendsen algorithm to control the temperature and pressure^54^. The time step was 2 femtoseconds (fs), and the SHAKE algorithm was used to constrain all bond lengths involving hydrogen atoms^55^. Long-range electrostatic interactions were treated using the particle mesh Ewald method^56^. Analysis of the trajectories was performed using the CPPTRAJ module of AmberTools16.

### Antibodies

Antibodies were purchased as follows: commercial mouse monoclonal antibodies to UL42 (13C9; Santa Cruz Biotechnology), glycoprotein B (gB) (H1817; Virusys), Flag (M2; Sigma) and Δ-actin (AC15; Sigma); rabbit polyclonal antibodies to human UNG (ab23926; Abcam;), APOBEC1 (A16756; ABclonal), green fluorescent protein (GFP) (598; MBL) and Flag (PM020; MBL); and rat monoclonal antibody to Hsc70 (1B5; Enzo Life Science). Mouse polyclonal antibody to vUNG and rabbit polyclonal antibodies to UL12 and ICP22 were reported previously^18, 57, 58^.

### Immunoblotting and immunofluorescence

Immunoblotting and immunofluorescence were performed as described previously^39, 59^. The amount of protein present in immunoblot bands was quantified using the ImageQuant LAS 4000 system with ImageQuant TL7.0 analysis software (GE Healthcare Life Sciences) or ChemiDoc MP (Bio-Rad) with ImageJ software according to the manufacturer’s instructions, and normalized to that of Δ-actin proteins.

### Production and purification of recombinant AAVs

HEK293EB cells were co-transfected with pAAV-ZsGreen1 (TaKaRa) or pAAV-UGI, along with pHelper (Addgene) and PHP.eB (Addgene) using PEI Max (Polysciences). At 10 days post-transfection, the culture supernatant was centrifuged at 10,000 ×g for 15 min at 4°C and filtrated through a 0.22-μm filter (Thermo Fisher Scientific). Then, recombinant AAVs were purified using an AAVpro Concentrator (TaKaRa) according to the manufacturer’s instructions, and designated AAV-ZsGreen and AAV-UGI. To measure the AAV titer, host genome and plasmid DNA were digested with DNase I (Sigma) before releasing the viral DNA from the particles, and then the viral DNA was released by Buffer AL (QIAGEN). The qPCR was performed using FastStart SYBR Green Master (Roche), a LightCycler 96 System (Roche) and primers against the inverted terminal repeats. The primer sequences were 5’-GGAACCCCTAGTGATGGAGTT- 3’ and 5’-CGGCCTCAGTGAGCGA-3’.

### Animal studies

For ocular HSV-1 infection, 4-week-old ICR mice were infected with 3×10^6^ plaque forming units (PFU)/eye of each of the indicated viruses as described previously^17–20^. Mice were monitored daily, and mortality from 1 to 21 days post-infection was attributed to the infected virus. For intracranial HSV-1 infection, 3-to-6-week-old C57BL/6 WT, APOBEC1-KO or APOBEC3-KO mice were infected intracranially with 1×10^3^ or 2×10^4^ PFU/head of the indicated viruses as described previously^18, 19, 22, 24^. Mice were monitored daily, and mortality from 1 to 21 days post-infection was attributed to the infected virus. Virus titers in the brains of mice were determined as described previously^18, 19, 22, 24^. For the administration of the recombinant AAV vector, 3-week-old C57BL/6 WT or APOBEC1- KO mice were infected intracranially with 5×10^10^ vg/head of AAV-ZsGreen or AAV-UGI using a 27-gauge needle (TOP) to penetrate the scalp and cranium over the hippocampal region of the left hemisphere with a needle guard to prevent penetration further than 3 mm. All animal experiments were carried out in accordance with the Guidelines for Proper Conduct of Animal Experiments, Science Council of Japan. The protocol was approved by the Institutional Animal Care and Use Committee (IACUC) of the Institute of Medical Science, The University of Tokyo (IACUC protocol approval numbers: PM27-113, PM27-73, PA17-67, PH2-12, A21-55, and A22-27).

### Quantitative reverse transcription PCR (qPCR)

Brains of ICR mice mock-infected or infected ocularly with 3×10^6^ PFU/eye of vUNG-S302A and vUNG-SA-repair were homogenized in TriPure isolation reagent (Roche) using a disposable pestle system (Fisher), and total RNA was then isolated with a High Pure RNA tissue kit (Roche) according to the manufacturer’s instructions. cDNA was synthesized from the isolated RNA with a Transcriptor First Strand cDNA synthesis kit (Roche) according to the manufacturer’s instructions. The amount of cDNA of specific genes was quantitated using the Universal ProbeLibrary (Roche) with TaqMan Master (Roche) and the LightCycler 96 system (Roche) according to the manufacturer’s instructions. Gene-specific primers and universal probes were designed using ProbeFinder software (Roche). The primer and probe sequences for mAPOBEC1 were 5’- CAGCGGTGTGACTATCCAGA-3’, 5’-TTGGCCAATAAGCTTCGTTT-3’, and Universal ProbeLibrary probe 67; those for mAPOBEC2 were 5’- CTCAAGTACAATGTCACCTGGTATG-3’, 5’-GTTTTGAGAATCCGGTCAGC-3’, and Universal ProbeLibrary probe 19; those for mAPOBEC3 were 5’- TACCAGCTGGAGCAGTTCAA-3’, 5’-CTGCATGCTGTTTGCCTTT-3’, and Universal ProbeLibrary probe 27; those for mAID were 5’-GCGCCCAGATCCAAAGTAT-3’, 5’- GCCATTTGTAATGGAAAACGA-3’, and Universal ProbeLibrary probe 72; and those for 18S rRNA were 5’-GCAATTATTCCCCATGAACG-3’, 5’-GGGACTTAATCAACGCAAGC-3’, and Universal ProbeLibrary probe 48. The expressions of mAPOBEC1, mAPOBEC2, mAPOBEC3, and mAID mRNAs were normalized to the amount of 18S rRNA expression. The relative amount of each gene expression was calculated using the comparative cycle threshold (2−ΔΔCT) method^41^.

### Hypermutation analysis

The 3D-PCR procedure, a highly sensitive assay for detecting AT-rich DNA, was performed as described previously, with minor modifications^44^. Briefly, HEp-2/ΔhUNGs cells in 6-well plates were transfected with HSV-1 infectious genome clones of vUNG-S302A or vUNG-SA-repair in combination with pcDNA-EGFP-P2A- hAPOBEC1, pcDNA-EGFP-P2A and pFlag-CMV2 (Sigma) by electroporation using a NEPA21 Super Electroporator (NepaGene). The electroporation was performed under the following conditions: poring pulse: 125 V, 3-ms pulse width, 50-ms pulse interval, two pulses, 10% attenuation rate, +; transfer pulse: 20 V, 50-ms pulse width, 50-ms pulse interval, five pulses, 40% attenuation rate, +/-. Six days post-transfection, the cells were lysed in lysis buffer and proteinase K from a GeneJET Genomic DNA Purification Kit (Thermo Fisher Scientific) and treated with RNase A (Thermo Fisher Scientific) according to the manufacturer’s instructions. After brief sonication, DNA was extracted with phenol-chloroform twice and diethyl ether four times, and then genome DNA was precipitated with ethanol. For the 3D-PCR of HSV-1 Us3^48^, the first PCR was performed as follows: 95°C for 9 min, 40 cycles of 95°C for 45 s, 63°C for 30 s, and 72°C for 1 min, with a final elongation step at 72°C for 1 min using the primers 5’-CAAACTGCCGCTCCTTAAAA-3’ and 5’-TCTGGGTGGCTGCTGTCAAA-3’. The nested PCR was performed as follows: 96-83°C for 5 min, 40 cycles of 96-83°C for 60 s, 62°C for 30 s, and 72°C for 1 min, with a final elongation step of 72°C for 10 min using the primers 5’-AATGGCCTGTCGTAAGTTTT-3’ and 5’-CTATGGGGTAGTCCTGGTTT-3’.

To determine the hypermutation frequency, PCR fragments from the nested PCR were cloned into pGEM-T-easy vectors (Promega), and the indicated number of successful recombinant clones was randomly selected and sequenced using a 3500 Genetic Analyzer (Thermo Fisher Scientific).

### Establishment of stable HEp-2 cells with tetracycline-inducible APOBEC1 expression

HEp-2 cells were transduced with supernatants of Plat-GP cells cotransfected with pMDG^60^ and pRetroX-Tet3G (TaKaRa), selected with 4 mg/ml G418 (Wako), and further transduced with supernatants of Plat-GP cells cotransfected with pMDG, and pRetroX- TRE3G-hAPOBEC1-3xflag or pRetroX-TRE3G-mAPOBEC1-3xflag, respectively. For the tetracycline-inducible (TetON) expression of hAPOBEC1 and mAPOBEC1 tagged with 3x flag, after double selection with 4 μg/ml puromycin (Sigma) and 1 mg/ml G418, resistant cells were designated HEp-2/TetON-hAPOBEC1-3xflag and HEp-2/TetON-mAPOBEC1-3xflag.

### Statistical analysis

Differences in relative cell viability were analyzed statistically using an unpaired two-tailed Student’s *t*-test. Differences in the relative UNG and vUNG activities were analyzed statistically using an unpaired two-tailed Student’s *t*-test and one-way ANOVA followed by Tukey’s test. Differences in the colocalization coefficient ratio and relative expressions of mRNA and proteins were analyzed statistically by one-way ANOVA followed by Tukey’s test. Differences in viral yields were analyzed statistically using the unpaired two-tailed Student’s *t*-test, Welch’s *t*-test and one-way ANOVA followed by Tukey’s test. Differences in the mortality rates of infected mice were analyzed statistically by the Log- rank test, and for four or three comparison analyses, *P-*values were sequentially considered significant after Holm’s sequentially rejective Bonferroni multiple-comparison adjustment.

## ACKNOWLEDGEMENTS and FUNDING INFORMATION

We thank Risa Abe, to, Tohru Ikegami and Yui Muto for their excellent technical assistance. This study was supported by Grants for Scientific Research and Grant-in-Aid for Scientific Research (S) (20H05692) from the Japan Society for the Promotion of Science (JSPS), grants for Scientific Research on Innovative Areas (21H00338, 21H00417, 22H04803) and a grant for Transformative Research Areas (22H05584) from the Ministry of Education, Culture, Science, Sports and Technology of Japan, a PRESTO grant (JPMJPR22R5) from Japan Science and Technology Agency (JST), grants (JP20wm0125002, JP22fk0108640, JP22gm1610008, JP223fa627001) from the Japan Agency for Medical Research and Development (AMED), grants from the International Joint Research Project of the Institute of Medical Science, the University of Tokyo and grants from the Takeda Science Foundation, the Cell Science Research Foundation, the MSD Life Science Foundation and the Mitsubishi Foundation.

The authors declare no competing financial interests.

---

**Supplementary Figure 1. Characterization of HEp-2/ΔhUNGs cells and the 100 ns MD simulations of vUNG. a**, The targeted hUNG1 and hUNG2 mutation sequences and the parental sequence in HEp-2/ΔhUNGs cells are shown. **b**, Lysates of HEp-2 and HEp- 2/ΔhUNGs cells were analyzed by immunoblotting with antibodies to hUNG or Δ-actin. Digital images are representative of three independent experiments. **c**, Cell viability of HEp-2 and HEp-2/ΔhUNGs cells. Each value is the mean ± SEM of the results of three independent experiments and is expressed relative to the mean for HEp-2 cells, which was normalized to 100%. The statistical significance determined by an unpaired two-tailed Student’s *t*-test is indicated. n.s., not significant. **d**, Uracil excision activity of lysis buffer (lane 1), cell lysates from HEp-2 (lane 2) and HEp-2/ΔhUNGs cells (lane 3). The cell lysates prepared for the UNG assays were analyzed by immunoblotting with antibodies to hUNG or Δ-actin. **e**. The amounts of cleaved oligonucleotides in the experiment in (**d**) were quantitated. Each value is the mean ± SEM of three independent experiments and is expressed relative to that in HEp-2 cell (lane2), which was normalized to 100%. The statistical significance determined by an unpaired two-tailed Student’s *t*-test is indicated. ****; *p* < 0.0001. **f**, Root-mean-square-deviation (RMSD) values of the backbone atoms for the non-phosphorylated vUNG and vUNG phosphorylated at Ser-302. **g**, RMSF values of each residue of the non-phosphorylated vUNG and vUNG phosphorylated at Ser-302.

**Supplementary Figure 2. Genome structure of recombinant viruses constructed in this study and multiple sequence alignment of uracil-DNA glycosylases. a,** Line 1, wild-type HSV-1(F) genome; line 2, structure of UL1, UL2 (vUNG) or UL3 CDS; lines 3 to 7, recombinant viruses with a mutation(s) in vUNG; and lines 8 to 12, recombinant viruses with mutation(s) in vUNG and carrying Flag-tagged vUNG (Flag-vUNG). **b**, Amino acid sequence alignment of UL2 (vUNG) homologs from HSV-1(F), HSV-2(HG52), VZV(VariVax), HCMV(AD169), HHV-6A(AJ), HHV-6B(Z29), HHV-7(RK), EBV(Akata) and KSHV(GK18), and UNG genes encoded by humans and *Escherichia* coli. Amino acid residues of the Water-activating loop of HSV-1 vUNG are shown in a green square. The residues conserved in the sequences and mutated in HSV-1 vUNG are shaded gray and orange, respectively.

**Supplementary Figure 3. Phosphorylation of vUNG Ser-302 contributes to its proper localization. a**, **b**. Confocal microscope images of HEp-2 cells infected with wild-type HSV- 1(F), Flag-vUNG-S53A, Flag-vUNG-S302A, Flag-vUNG-SA-repair, Flag-vUNG- Q177L/D178N, or Flag-vUNG-QL/DN-repair for 24 h at an MOI of 10 and stained with antibodies to Flag (**a**, **b**), UL42 (vPOL processivity factor) (**a**) and Hsc-70 (**b**). Scale bar, 10 μm. Digital images are representative of three independent experiments. **c**, **f**. Colocalization in the experiments of (**a**) and (**b**) were quantified using Pearson’s colocalization coefficient (PCC). Each value is the mean ± SEM (*n* = 6). The statistical significance determined by one-way ANOVA followed by Tukey’s test is indicated. ***; *p* < 0.001. n.s., not significant. **d**, **e**, **g**, **h**. Line-scan analysis of colocalization in the experiments in (**a**) and (**b**). The fluorescence intensities of white arrows in the panels for Flag-vUNG-S302A and Flag-vUNG- Q177L/D178N were determined.

**Supplementary Figure 4. Characterization of a recombinant virus constructed in this study. a** to **f**, **i**, and **j**, Vero cells were infected with HSV-1(F) (**a** to **f**, **i** and **j**), vUNG-S53A (**a**, **b**), vUNG-S302A (**c**, **d**), vUNG-SA-repair (**c**, **d**), vUNG-Q177L/D178N (**e**, **f**), vUNG-QL/DN-repair (**e**, **f**), Flag-vUNG-S53A (**i**, **j**), Flag-vUNG-S302A (**i**, **j**), Flag-vUNG-SA-repair (**i**, **j**), Flag-vUNG-Q177L/D178N (**i**, **j**), or Flag-vUNG-QL/DN-repair (**i**, **j**) at an MOI of 10 (**a**, **c**, **e, i**) or 0.01 (**b**, **d**, **f, j**). Total virus from the cell culture supernatants and infected cells was harvested at the indicated times and assayed. Each value represents the mean ± SEM of three independent experiments. The statistical significance determined by one-way ANOVA followed by Tukey’s test is indicated. n.s., not significant. **g**, **h** and **k**, Vero cells mock-infected (**g**, **h**, **k**) or infected with wild-type HSV-1(F) (**g**, **h**, **k**), vUNG-S53A (**g**), vUNG-S302A (**g**), vUNG-SA-repair (**g**), vUNG-Q177L/D178N (**h**), vUNG-QL/DN-repair (**h**), Flag-vUNG-S53A (**k**), Flag-vUNG-S302A (**k**), Flag-vUNG-SA-repair (**k**), Flag-vUNG-Q177L/D178N (**k**), or Flag-vUNG-QL/DN-repair (**k**) for 24 h at an MOI of 10 were lysed and analyzed by immunoblotting with antibodies to vUNG (**g**, **h**, **k**), Flag (**k**), UL12 (**g**, **h**, **k**) and β-actin (**g**, **h**, **k**). Digital images are representative of three independent experiments.

**Supplementary Figure 5. Viral replication of HSV-1(F) and vUNG mutant viruses in HEp-2/ΔhUNGs cells. a** to **d**, Viral titers of HEp-2/ΔhUNGs were infected with wild-type HSV-1(F) (**a** to **d**), vUNG-S302A (**a**, **b**) or vUNG-Q177L/D178N (**c**, **d**) for 24 h at an MOI of 10 (**a**, **c**) or for 48 h at an MOI of 0.01 (**b**, **d**), respectively, and assayed. Each value is the mean ± SEM of four independent experiments. The statistical significance determined by an unpaired two-tailed Student’s *t*-test is indicated. n.s., not significant.

**Supplementary Figure 6. Effects of mutations in vUNG and/or knock-out of APOBEC1 on the mortality of mice following intracranial infection. a**, **c**, **e**, Four-week-old C57BL/6 mice were infected intracranially with 1×10^3^ PFU/head of vUNG-S302A (**a**, *n* = 16; **e**, *n* = 13), vUNG-SA-repair (**a**, *n* = 16), vUNG-Q177L/D178N (**c**, *n* = 9; **e**, *n* = 12), or vUNG-QL/DN-repair (**c**, *n* = 8) and monitored daily for survival for 21 days. The statistical significance determined by Log-rank test is indicated. *; *p* < 0.05, n.s. not significant. **b**, **d**, **f**, Five-week-old C57BL/6 mice were infected intracranially with 1×10^3^ PFU/head of vUNG-S302A (**b**, *n* = 16; **f**, *n* = 9), vUNG-SA-repair (**b**, *n* = 15), vUNG-Q177L/D178N (**d**, *n* = 12; **f**, *n* = 10), or vUNG-QL/DN-repair (**d**, *n* = 9). Viral titers in the brains of infected mice at 5 days post-infection were assayed. Dashed lines indicate the limit of detection. n.d. not detected. Each value is the mean ± SEM for each group. The statistical significance determined by Welch’s *t*-test is indicated. *; *p* < 0.05, n.s. not significant. **g**, Three-to-six-week-old C57BL/6 WT, or APOBEC1-KO mice were infected intracranially with 1×10^3^ PFU/head of vUNG- Q177L/D178N (n = 16), and monitored daily for 21 days. The statistical significance determined by Log-rank test is indicated. *; *p* < 0.05. **h**, Three-to-six-week-old C57BL/6 WT (*n* = 13) or APOBEC1-KO (*n* = 9) mice were infected intracranially with 1×10^3^ PFU/head of vUNG-Q177L/D178N. Viral titers in the brains of infected mice at 5 days post-infection were assayed. Each value is the mean ± SEM for each group. The statistical significance determined by Welch’s *t*-test is indicated. *; *p* < 0.05, n.s. not significant.

**Supplementary Figure 7. Results of 3D-PCR at a denaturation temperature of 93.8°C, localization of APOBEC1 in HSV-1-infected HEp-2 cells**, **and robust transduction of the brain by AAV vectors. a** to **c**, In the experiment in Fig. 5a, Us3 DNA amplified at a denaturation temperature of 93.8°C (**a**: lane 1, **b**: lane 4 and **c**: lane 7 in Fig. 5a) was excised, cloned, and sequenced. Results are summarized: C-to-T and G-to-A hypermutations are indicated as blue and orange vertical lines, respectively. All other base substitutions are shown as green vertical lines. **d**, Parental HEp-2, HEp-2/TetON-hAPOBEC1-3xflag and HEp- 2/TetON-mAPOBEC1-3xflag cells were mock treated or treated with doxycycline (DOX) (1 μg/ml), harvested at 72 h post-treatment, and analyzed by immunoblotting with antibodies to APOBEC1, Flag and Δ-actin. Digital images are representative of three independent experiments. **e**, Confocal microscope images of parental HEp-2, HEp-2/TetON-hAPOBEC1-3xflag and HEp-2/TetON-mAPOBEC1-3xflag cells infected with wild-type HSV-1(F) for 9 h at an MOI of 10 in the presence of DOX and stained with antibody to Flag and ICP22. Digital images are representative of three independent experiments. **f**, Three-week-old C57BL/6 WT or APOBEC1-KO mice were mock-treated (*n* = 2) or treated pretreated intracranially with 5×10^10^ vg/head of AAV-ZsGreen (*n* = 3), respectively. At 14 days after pretreatment with AAV-vectors, mice were pretreated intracranially with 2×10^4^ PFU/head of vUNG-SA-repair. Bright field and fluorescent whole-brain images were obtained by using ChemiDoc™ Touch MP at 4 days post-infection with HSV-1. Digital images are representative of each group.

**Supplementary Figure 8. Expression of hAPOBEC1 in the human brain. a** to **c**, Images taken from the Allen Human Brain Atlas (https://human.brain-map.org/) show the expression of hAPOBEC1 in the left- (**a**), right- (**b**) and sub- (**c**) cortex of an adult human brain, respectively. The database contains data of postmortem brains from men and women between the ages of 18 and 68 years with no known neuropsychiatric or neuropathological history.

**Supplementary Figures 9 to 10. Uncropped images of immunoblots, autoradiography and agarose gel stained with ethidium bromide in this study.** Blue boxes indicate the cropped areas shown in the indicated figures. Molecular weight is indicated as a number.

