## Supplementary Figures 1-8, and Supplementary Table 1-4 for "Evasion of APOBEC1-mediated Intrinsic Immunity by a Herpesvirus Uracil DNA Glycosylase Is a Determinant of Viral Encephalitis"

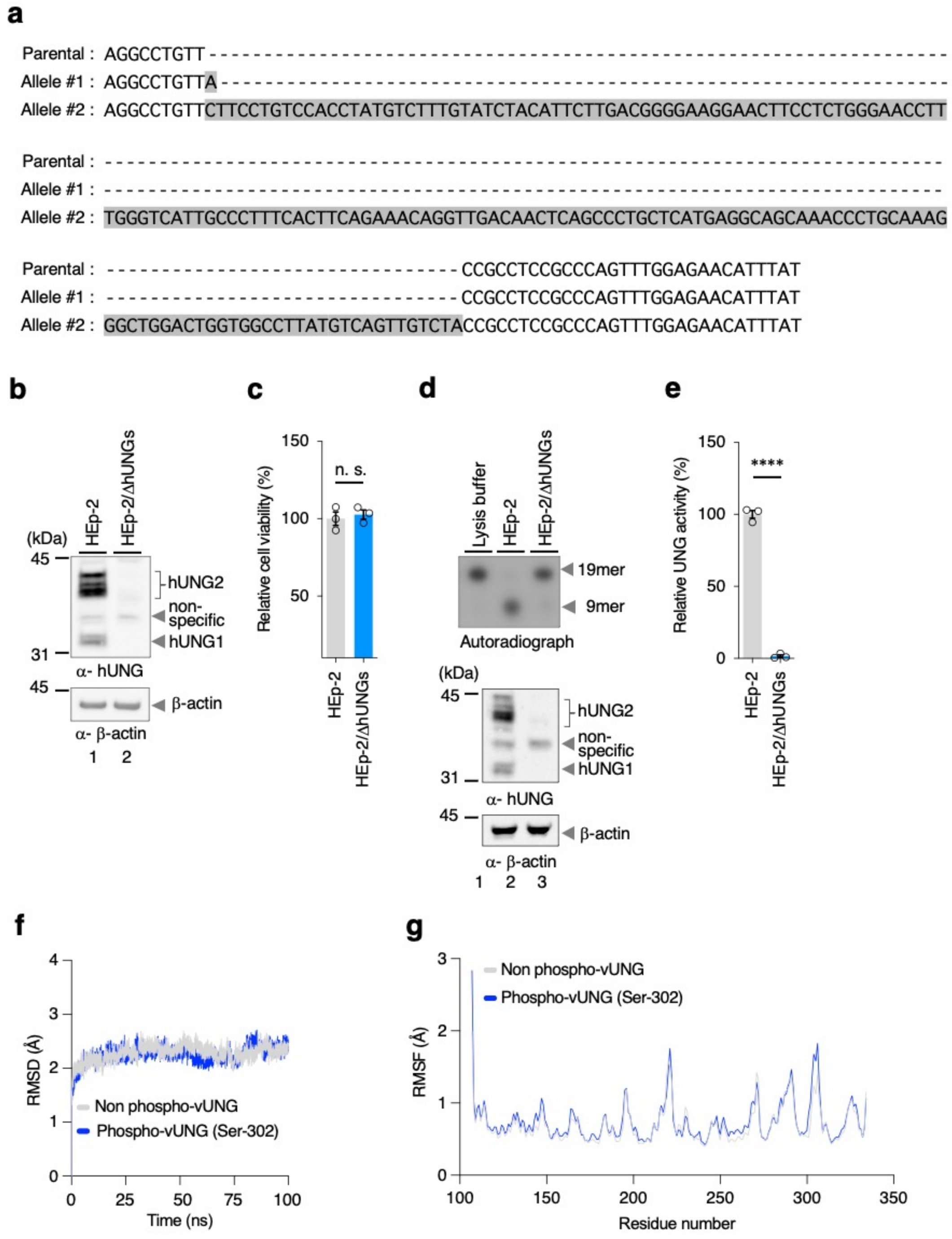

**Supplementary FIG.1 A. Kato and H. Harima et al**

**Supplementary Figure 1. Characterization of HEp-2/ΔhUNGs cells and the 100 ns MD simulations of vUNG.** **a**, The targeted hUNG1 and hUNG2 mutation sequences and the parental sequence in HEp-2/ΔhUNGs cells are shown. **b**, Lysates of HEp-2 and HEp-2/ΔhUNGs cells were analyzed by immunoblotting with antibodies to hUNG or β-actin. Digital images are representative of three independent experiments. **c**, Cell viability of HEp-2 and HEp-2/ΔhUNGs cells. Each value is the mean ± SEM of the results of three independent experiments and is expressed relative to the mean for HEp-2 cells, which was normalized to 100%. The statistical significance determined by an unpaired two-tailed Student's *t*-test is indicated. n.s., not significant. **d**, Uracil excision activity of lysis buffer (lane 1), cell lysates from HEp-2 (lane 2) and HEp-2/ΔhUNGs cells (lane 3). The cell lysates prepared for the UNG assays were analyzed by immunoblotting with antibodies to hUNG or β-actin. **e**. The amounts of cleaved oligonucleotides in the experiment in (**d**) were quantitated. Each value is the mean ± SEM of three independent experiments and is expressed relative to that in HEp-2 cell (lane2), which was normalized to 100%. The statistical significance determined by an unpaired two-tailed Student's *t*-test is indicated. ****; $p < 0.0001$. **f**, Root-mean-square-deviation (RMSD) values of the backbone atoms for the non-phosphorylated vUNG and vUNG phosphorylated at Ser-302. **g**, RMSF values of each residue of the non-phosphorylated vUNG and vUNG phosphorylated at Ser-302.

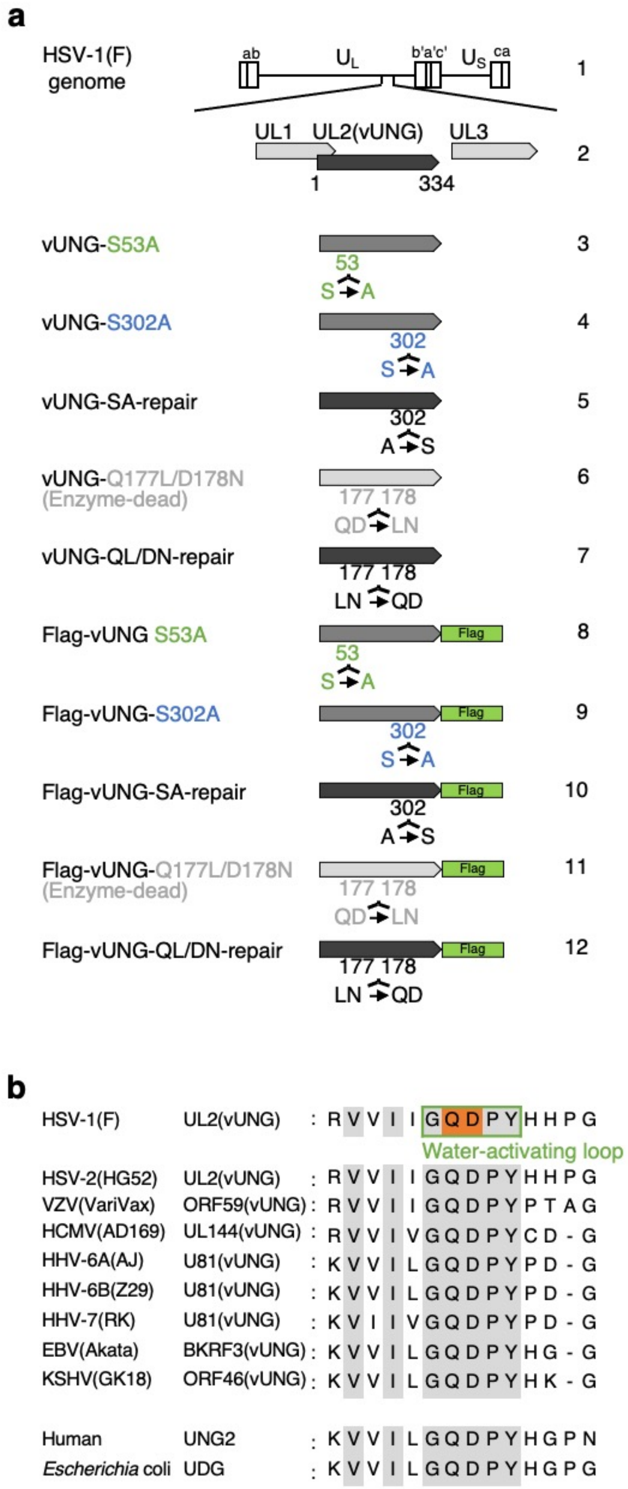

**Supplementary FIG.2 A. Kato and H. Harima et al**

**Supplementary Figure 2. Genome structure of recombinant viruses constructed in this study and multiple sequence alignment of uracil-DNA glycosylases.** **a**, Line 1, wild-type HSV-1(F) genome; line 2, structure of UL1, UL2 (vUNG) or UL3 CDS; lines 3 to 7, recombinant viruses with a mutation(s) in vUNG; and lines 8 to 12, recombinant viruses with mutation(s) in vUNG and carrying Flag-tagged vUNG (Flag-vUNG). **b**, Amino acid sequence alignment of UL2 (vUNG) homologs from HSV-1(F), HSV-2(HG52), VZV(VariVax), HCMV(AD169), HHV-6A(AJ), HHV-6B(Z29), HHV-7(RK), EBV(Akata) and KSHV(GK18), and UNG genes encoded by humans and Escherichia coli. Amino acid residues of the Water-activating loop of HSV-1 vUNG are shown in a green square. The residues conserved in the sequences and mutated in HSV-1 vUNG are shaded gray and orange, respectively.

.

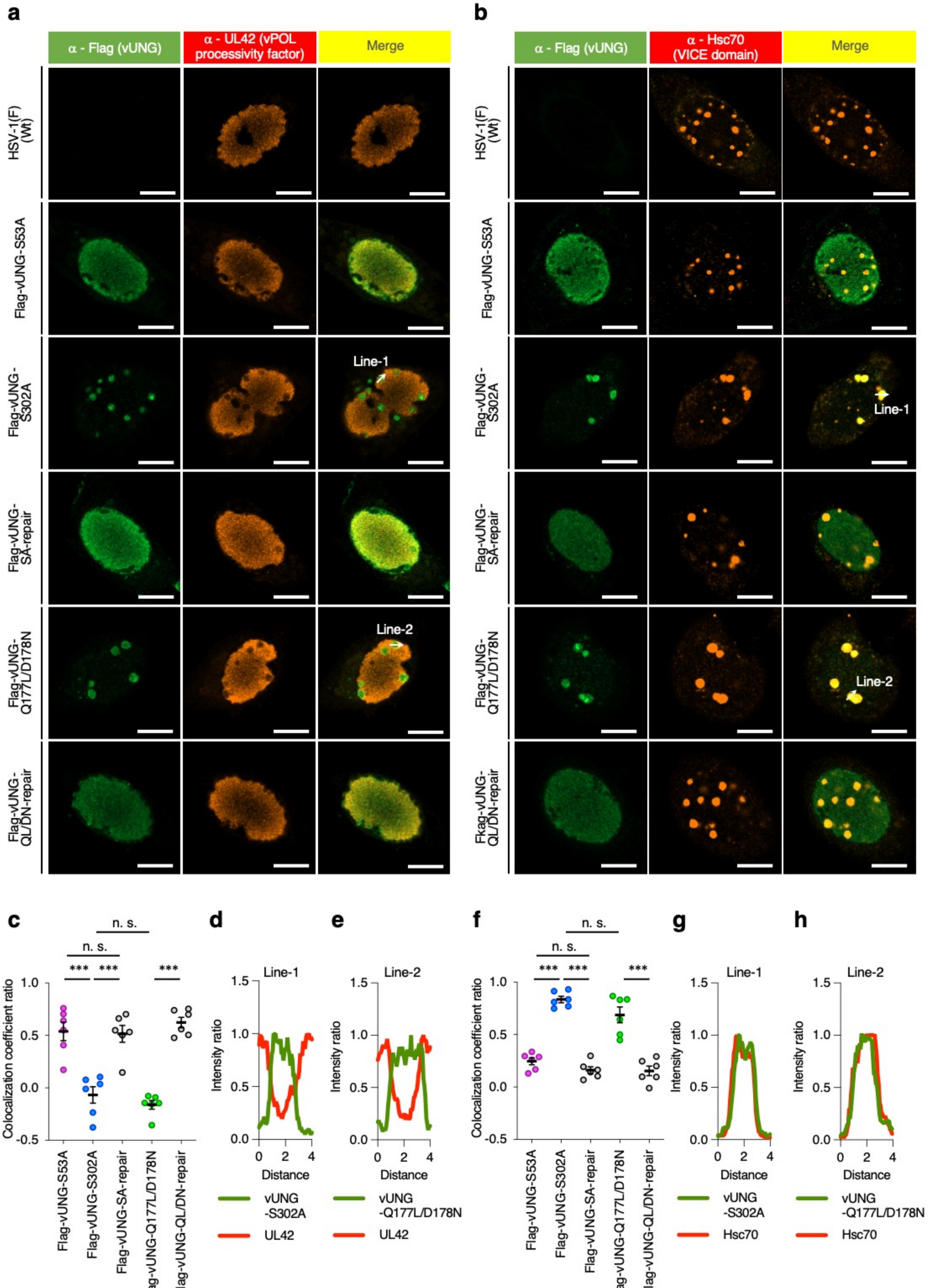

**Supplementary FIG.3 A. Kato and H. Harima et al**

**Supplementary Figure 3. Phosphorylation of vUNG Ser-302 contributes to its proper localization. a, b**. Confocal microscope images of HEp-2 cells infected with wild-type HSV-1(F), Flag-vUNG-S53A, Flag-vUNG-S302A, Flag-vUNG-SA-repair, Flag-vUNG-Q177L/D178N, or Flag-vUNG-QL/DN-repair for 24 h at an MOI of 10 and stained with antibodies to Flag (**a**, **b**), UL42 (vPOL processivity factor) (**a**) and Hsc-70 (**b**). Scale bar, 10 µm. Digital images are representative of three independent experiments. **c**, **f**. Colocalization in the experiments of (**a**) and (**b**) were quantified using Pearson's colocalization coefficient (PCC). Each value is the mean ± SEM (n = 6). The statistical significance determined by one-way ANOVA followed by Tukey's test is indicated. ***; $p < 0.001$. n.s., not significant. **d**, **e**, **g**, **h**. Line-scan analysis of colocalization in the experiments in (**a**) and (**b**). The fluorescence intensities of white arrows in the panels for Flag-vUNG-S302A and Flag-vUNG-Q177L/D178N were determined.

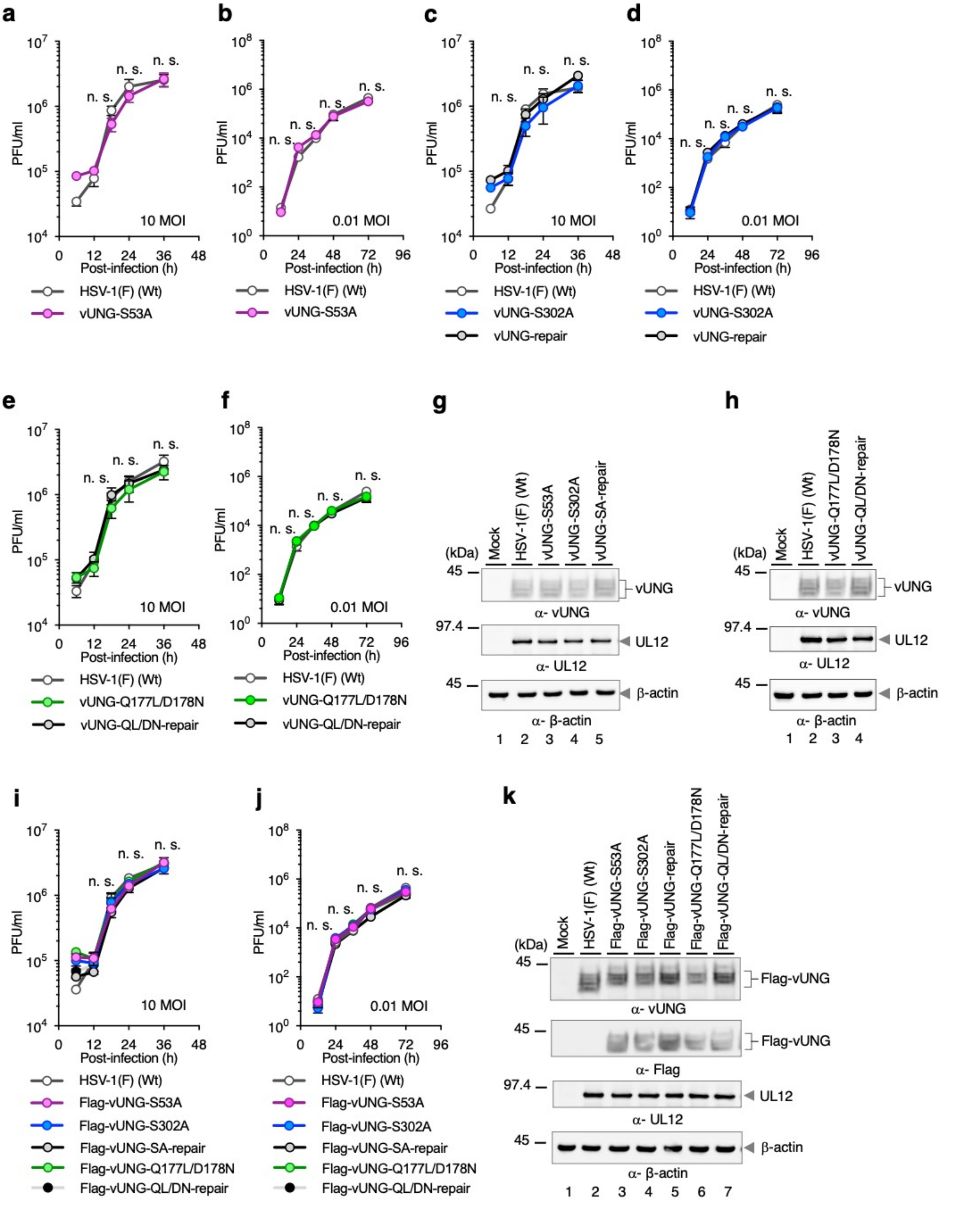

**Supplementary FIG.4 A. Kato and H. Harima et al**

**Supplementary Figure 4. Characterization of a recombinant virus constructed in this study. a** to **f**, **i**, and **j**, Vero cells were infected with HSV-1(F) (**a** to **f**, **i** and **j**), vUNG-S53A (**a**, **b**), vUNG-S302A (**c**, **d**), vUNG-SA-repair (**c**, **d**), vUNG-Q177L/D178N (**e**, **f**), vUNG-QL/DN-repair (**e**, **f**), Flag-vUNG-S53A (**i**, **j**), Flag-vUNG-S302A (**i**, **j**), Flag-vUNG-SA-repair (**i**, **j**), Flag-vUNG-Q177L/D178N (**i**, **j**), or Flag-vUNG-QL/DN-repair (**i**, **j**) at an MOI of 10 (**a**, **c**, **e**, **i**) or 0.01 (**b**, **d**, **f**, **j**). Total virus from the cell culture supernatants and infected cells was harvested at the indicated times and assayed. Each value represents the mean ± SEM of three independent experiments. The statistical significance determined by one-way ANOVA followed by Tukey's test is indicated. n.s., not significant. **g**, **h** and **k**, Vero cells mock-infected (**g**, **h**, **k**) or infected with wild-type HSV-1(F) (**g**, **h**, **k**), vUNG-S53A (**g**), vUNG-S302A (**g**), vUNG-SA-repair (**g**), vUNG-Q177L/D178N (**h**), vUNG-QL/DN-repair (**h**), Flag-vUNG-S53A (**k**), Flag-vUNG-S302A (**k**), Flag-vUNG-SA-repair (**k**), Flag-vUNG-Q177L/D178N (**k**), or Flag-vUNG-QL/DN-repair (**k**) for 24 h at an MOI of 10 were lysed and analyzed by immunoblotting with antibodies to vUNG (**g**, **h**, **k**), Flag (**k**), UL12 (**g**, **h**, **k**) and β-actin (**g**, **h**, **k**). Digital images are representative of three independent experiments.

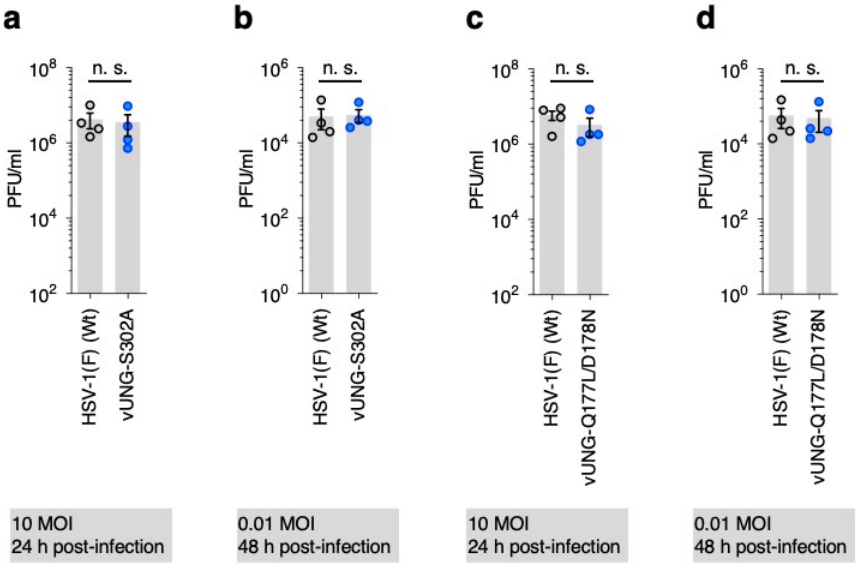

**Supplementary Figure 5. Viral replication of HSV-1(F) and vUNG mutant viruses in HEp-2/ΔhUNGs cells. a** to **d**, Viral titers of HEp-2/ΔhUNGs were infected with wild-type HSV-1(F) (**a** to **d**), vUNG-S302A (**a**, **b**) or vUNG-Q177L/D178N (**c**, **d**) for 24 h at an MOI of 10 (**a**, **c**) or for 48 h at an MOI of 0.01 (**b**, **d**), respectively, and assayed. Each value is the mean ± SEM of four independent experiments. The statistical significance determined by an unpaired two-tailed Student's *t*-test is indicated. n.s., not significant.

**Supplementary FIG.5 A. Kato and H. Harima et al**

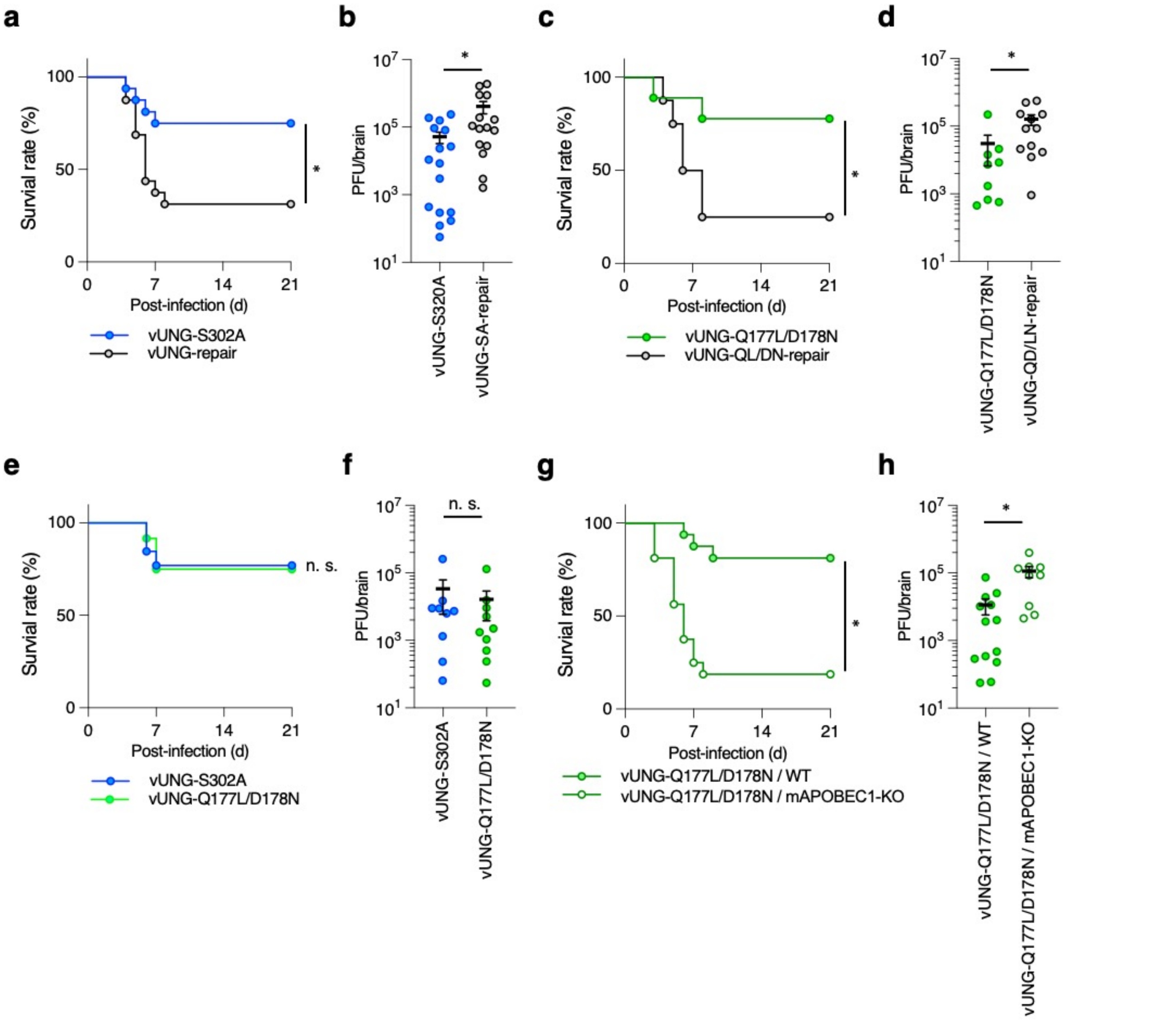

**Supplementary FIG.6 A. Kato and H. Harima et al**

**Supplementary Figure 6. Effects of mutations in vUNG and/or knock-out of APOBEC1 on the mortality of mice following intracranial infection.** **a**, **c**, **e**, Four-week-old C57BL/6 mice were infected intracranially with $1 \times 10^3$ PFU/head of vUNG-S302A (**a**, n = 16; **e**, n = 13), vUNG-SA-repair (**a**, n = 16), vUNG-Q177L/D178N (**c**, n = 9; **e**, n = 12), or vUNG-QL/DN-repair (**c**, n = 8) and monitored daily for survival for 21 days. The statistical significance determined by Log-rank test is indicated. *; $p < 0.05$, n.s. not significant. **b**, **d**, **f**, Five-week-old C57BL/6 mice were infected intracranially with $1 \times 10^3$ PFU/head of vUNG-S302A (**b**, n = 16; f, n = 9), vUNG-SA-repair (**b**, n = 15), vUNG-Q177L/D178N (**d**, n = 12; f, n = 10), or vUNG-QL/DN-repair (**d**, n = 9). Viral titers in the brains of infected mice at 5 days post-infection were assayed. Dashed lines indicate the limit of detection. n.d. not detected. Each value is the mean ± SEM for each group. The statistical significance determined by Welch's *t*-test is indicated. *; $p < 0.05$, n.s. not significant. **g**, Three-to-six-week-old C57BL/6 WT, or APOBEC1-KO mice were infected intracranially with $1 \times 10^3$ PFU/head of vUNG-Q177L/D178N (n = 16), and monitored daily for 21 days. The statistical significance determined by Log-rank test is indicated. *; $p < 0.05$. **h**, Three-to-six-week-old C57BL/6 WT (n = 13) or APOBEC1-KO (n = 9) mice were infected intracranially with $1 \times 10^3$ PFU/head of vUNG-Q177L/D178N. Viral titers in the brains of infected mice at 5 days post-infection were assayed. Each value is the mean ± SEM for each group. The statistical significance determined by Welch's *t*-test is indicated. *; $p < 0.05$, n.s. not significant.

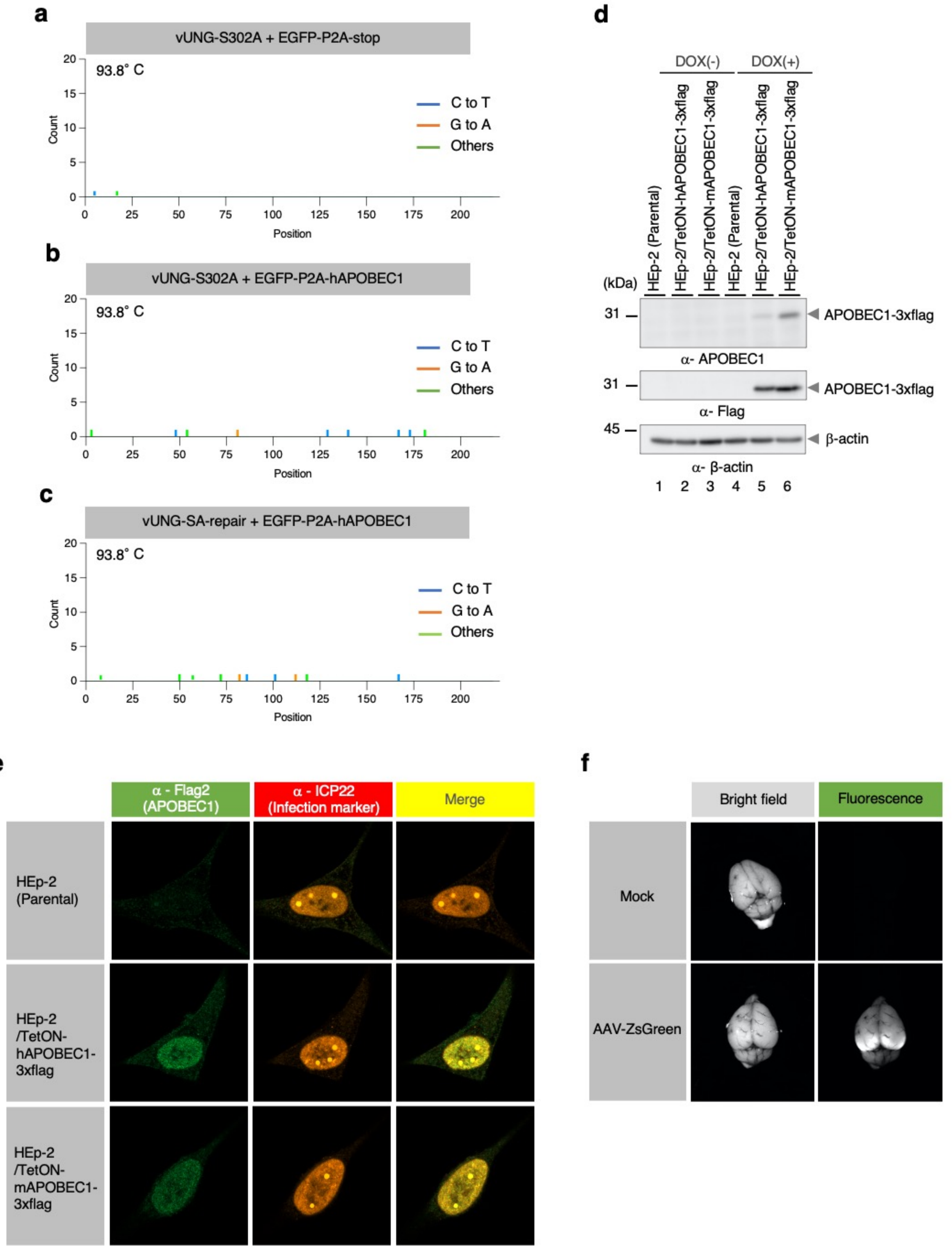

**Supplementary FIG.7 A. Kato and H. Harima et al**

**Supplementary Figure 7. Results of 3D-PCR at a denaturation temperature of 93.8°C, localization of APOBEC1 in HSV-1-infected HEp-2 cells, and robust transduction of the brain by AAV vectors. a** to **c**, In the experiment in Fig. 5**a**, Us3 DNA amplified at a denaturation temperature of 93.8°C (**a**: lane 1, **b**: lane 4 and **c**: lane 7) was excised, cloned, and sequenced. Results are summarized: C-to-T and G-to-A hypermutations are indicated as blue and orange vertical lines, respectively. All other base substitutions are shown as green vertical lines. **d**, Parental HEp-2, HEp-2/TetON-hAPOBEC1-3xflag and HEp-2/TetON-mAPOBEC1-3xflag cells were mock treated or treated with doxycycline (DOX) (1 μg/ml), harvested at 72 h post-treatment, and analyzed by immunoblotting with antibodies to APOBEC1, Flag and β-actin. Digital images are representative of three independent experiments. **e**, Confocal microscope images of parental HEp-2, HEp-2/TetON-hAPOBEC1-3xflag and HEp-2/TetON-mAPOBEC1-3xflag cells infected with wild-type HSV-1(F) for 9 h at an MOI of 10 in the presence of DOX and stained with antibody to Flag and ICP22. Digital images are representative of three independent experiments. **f**, Three-week-old C57BL/6 WT or APOBEC1-KO mice were mock-treated (n = 2) or treated pretreated intracranially with $5 \times 10^{10}$ vg/head of AAV-ZsGreen (n = 3), respectively. At 14 days after pretreatment with AAV-vectors, mice were pretreated intracranially with $2 \times 10^{4}$ PFU/head of vUNG-SA-repair. Bright field and fluorescent whole-brain images were obtained by using ChemiDoc™ Touch MP at 4 days post-infection with HSV-1. Digital images are representative of each group.

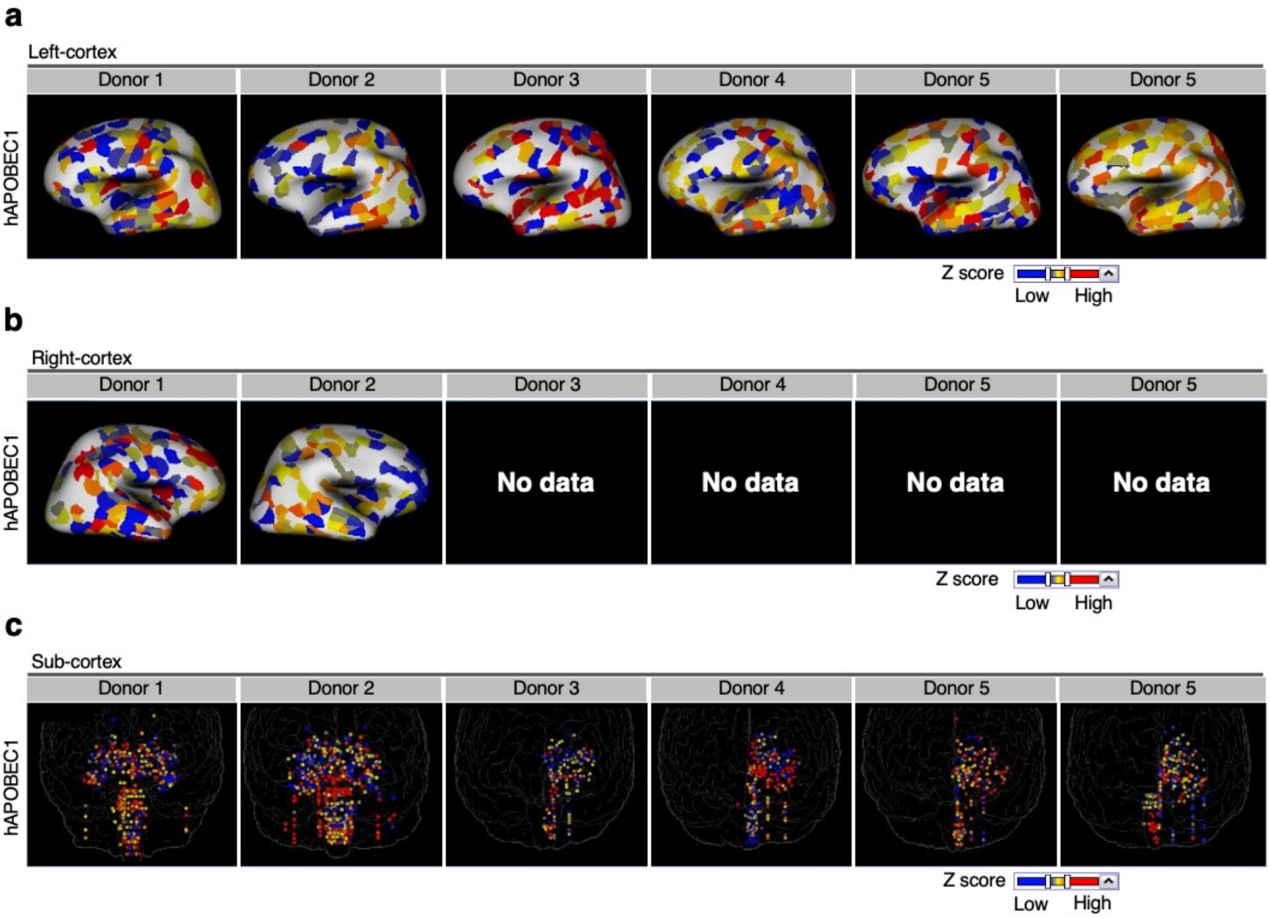

**Supplementary Figure 8. Expression of hAPOBEC1 in the human brain. a** to **c**, Images taken from the Allen Human Brain Atlas (https://human.brain-map.org/) show the expression of hAPOBEC1 in the left- (**a**), right- (**b**) and sub- (**c**) cortex of an adult human brain, respectively. The database contains data of postmortem brains from men and women between the ages of 18 and 68 years with no known neuropsychiatric or neuropathological history.

**Supplementary Table 1. vUNG phosphopeptides and phosphorylation sites identified by MS/MS.**

| Phosphorylation site(s)[a] | Phosphopeptide sequence[b] | Highest Score[c] |
|---|---|---|
| Thr-38 and Ser-53 | ADADDP(p)TPGASNDAATETRPG(p)SGGEPAACR | 70.16 |
| Thr-47 | ADADDPTPGASNDAA(p)TETRPGSGGEPAACR | 47.83 |
| Ser-328 | SI(p)SPIDWSV | 31.35 |
| Ser-302 | FSHP(p)SPLSK | 29.24 |
| Ser-16 and Thr-21 | RP(p)SSPRW(p)TPPR | 25.52 |
| Ser-6 and Ser-8 | AC(p)SR(p)SPSPR | 24.61 |
| Thr-21 and Thr-27 | W(p)TPPRDG(p)TPPQK | 23.29 |
| Thr-21 | W(p)TPPRDGTPPQK | 22.23 |
| Ser-6 and Ser-8 | RAC(p)SR(p)SPSPR | 22.90 |
| Ser-16 and Ser-17 | RP(p)S(p)SPRWTPPR | 22.02 |

[a]Evaluation of phosphorylation sites was conducted on the basis of a threshold of above 75% phosphoRS site probabilities, which were calculated by Proteome Discoverer (ver. 1.3; Thermo Fisher Scientific).

[b]In the peptide sequences, (p) indicates a phosphorylated amino acid.

[c]Highest Score represents the highest mascot ion score of the corresponding peptide(s).

### Supplementary Table 2. Detailed data of mutations in 3D-PCR analyses.

#### 88.9°C_vUNG-S302A + EGFP-P2A-hAPOBEC1

| Clone number | Amino acid sequences of wild-type Us3 | | | | | | | | | | | | | | | | | | | | | | | | | | | | | | | | | | | | | | | | |
|---|---|---|---|---|---|---|---|---|---|---|---|---|---|---|---|---|---|---|---|---|---|---|---|---|---|---|---|---|---|---|---|---|---|---|---|---|---|---|---|---|---|
| | g | t | c | g | c | g | t | t | t | a | c | g | g | g | g | g | a | c | a | g | g | g | c | a | g | g | a | g | g | a | a | g | g | a | g | g | a | g | g | c | c | g |
| 1 | | | T | | | | | | | | | | | | | | | | | | | | | | | | | | | | | | | | | | | | | | T | |
| 2 | | | T | | | | | | | | | | | | | | | | | | | | | | | | | | | | | | | | | | | | | | | |
| 3 | | | T | | | | | | | | | | | | | | | | | | | | | | | | | | | | | | | | | | | | | | | |
| 4 | | | | | | | | | | | | | | | | | | | | | | | | | | | | | | | | | | | | | | | | | T | |
| 5 | | | | | | | | | | | | | | | | | | | | | | | | | | | | | | | | | | | | | | | | | T | |
| 6 | | | T | | | | | | | | | | | | | | | | | | | | | | | | | | | | | | | | | | | | | | | |
| 7 | | | T | A | | | | | | | | | | | | | | | | | | | | | | | | | | | | | | | | | | | | | | |
| 8 | | | T | | | | | | | | | | | | | | | | | | | | | | | | | | | | | | | | | | | | | | | |
| 9 | | | | | | | | | | | | | | | | | | | | | | | | | | | | | | | | | | | | | | | | | | |
| 10 | | | | | | | | | | | | | | | | | | | | | | | | | | | | | | | | | | | | | | | | | | |
| 11 | | | T | | | | | | | | | | | | | | | | | | | | | | | | | | | | | | | | | | | | | | | |
| 12 | | | | | | | | | | | | | | | | | | | | | | | | | | | | | | | | | | | | | | | | | T | |
| 13 | | | T | | | | | | | | | | | | | | | | | | | | | | | | | | | | | | | | | | | | | | | |
| 14 | | | T | | | | | | | | | | | | | | | | | | | | | | | | | | | | | | | | | | | | | | | |
| 15 | | | T | | | | | | | | | | | | | | | | | | | | | | | | | | | | | | | | | | | | | | | |
| 16 | | | | | | | | | | | | | | | | | | | | | | | | | | | | | | | | | | | | | | | | | | |
| 17 | | | T | | | | | | | | | | | | | | | | | | | T | | | | | | | | | | | | | | | | | | | | |
| 18 | | | T | | | | | | | | | | | | | | | | | | | | T | | | | | | | | | | | | | | | | | | | |
| 19 | | | T | | | | | | | | | | | | | | | | | | | | | | | | | | | | | | | | | | | | | | | |
| 20 | | | T | | | | | | | | | | | | | | | T | | | | | | | | | | | | | | | | | | | | | | T | | |
| C to T | 0 | 0 | 14 | 0 | 0 | 0 | 0 | 0 | 0 | 0 | 0 | 0 | 0 | 0 | 0 | 0 | 0 | 1 | 0 | 0 | 0 | 0 | 1 | 0 | 0 | 0 | 0 | 0 | 0 | 0 | 0 | 0 | 0 | 0 | 0 | 0 | 0 | 0 | 0 | 1 | 4 | 0 |
| G to A | 0 | 0 | 0 | 0 | 0 | 0 | 0 | 0 | 0 | 0 | 0 | 0 | 0 | 0 | 0 | 0 | 0 | 0 | 0 | 0 | 0 | 0 | 0 | 0 | 0 | 0 | 0 | 0 | 0 | 0 | 0 | 0 | 0 | 0 | 0 | 0 | 0 | 0 | 0 | 0 | 0 | 0 |
| Other | 0 | 0 | 0 | 1 | 0 | 0 | 0 | 0 | 0 | 0 | 0 | 0 | 0 | 0 | 0 | 0 | 0 | 0 | 0 | 0 | 0 | 1 | 0 | 0 | 0 | 0 | 0 | 0 | 0 | 0 | 0 | 0 | 0 | 0 | 0 | 0 | 0 | 0 | 0 | 0 | 0 | 0 |

| Clone number | t | c | c | c | g | c | c | g | g | a | g | a | c | a | a | a | g | c | c | g | t | c | c | c | g | g | g | t | g | t | t | t | c | c | t | c | a | t | g | g | c | c | c | c | t |
|---|---|---|---|---|---|---|---|---|---|---|---|---|---|---|---|---|---|---|---|---|---|---|---|---|---|---|---|---|---|---|---|---|---|---|---|---|---|---|---|---|---|---|---|---|---|
| 1 | | T | T | T | | | T | | | | | | | | | | | | T | | | T | T | T | | | | | | | | | T | T | | T | | | | | T | T | T | T | |
| 2 | | | | | | | | | | | | | | | | | | | | | | T | | | | | | | | | | | T | | | T | | | | | | | T | | |
| 3 | | | | | | | | | | | | | | | | | | | | | | T | | | | | | | | C | | | T | | | T | | | | | | | T | | |
| 4 | | | | | | | | | | | | | | | | | | | | | | | | | | | | | | | | | T | | | | | | | | | T | T | T | |
| 5 | | | | | | | | | | | | | | | | | | | | | | | | G | | | | | | | | | T | | | | | | | | | T | T | T | |
| 6 | | | | | | | | | | | | | | | | | | | | | | T | | | | | | | | | | | T | | | T | | | | | | | T | | |
| 7 | | | | | | | | | | | | | | | | | | | | | | T | | | | | | | | | | | T | | | T | | | | | | | T | | |
| 8 | | | | | | | | | | | | | | | | | | | | | | T | | | | | | | | | | | T | | | T | | | | | | | T | | |
| 9 | | | | | | | | | | | | T | | | | | | | | | | T | | | | | | | | | | | T | | | T | | | | | | | T | | |
| 10 | | | | | | | | | | | | | | | | | | | | | | T | | | | | | | | | | | T | | | T | | | | | | | T | | |
| 11 | | | | | | | | | | | | | | | | | | | | | | T | | | | | | | | | | | T | | | T | | | | | | | T | | |
| 12 | | T | T | T | | | | | | | | | | | | | | | | | | T | T | T | | | | | | | | | T | T | | T | | | | | | T | T | T | |
| 13 | | T | | | | | | | | | | | | | | | | | | | | | | | | | | | | | | | T | | | T | | | | | | | T | | |
| 14 | | T | | | | | | | | | | | | | | | | | | | | | | | | | | | | | | | T | | | T | | | | | | | T | | |
| 15 | | | | | | | | | | | | | | | | | | | | | | T | | | | | | | | | | | T | | | T | | | | | | | T | | |
| 16 | | T | T | T | | T | T | | | | | | T | | | | | | T | | | | | | | | | | | | | | | | | | | | | | | | | T | |
| 17 | | | | | | | | | | | | | | | | | | | | | | T | | | | | | | | | | | T | | | T | | | | | | | T | | |
| 18 | | | | | | | | | | | | | | | | | | | | | | T | | | | | | | | | | | T | | | T | | | | | | | T | | |
| 19 | | | | | | | | | | | | | | | | | | | | | | T | | | | | | | | | | | T | | | T | | | | | | | T | | |
| 20 | | | | | | | | | | | | | | | | | | | | | | T | | | | | | | | | | | T | | | T | | | | | | | T | | |
| C to T | 0 | 5 | 3 | 3 | 0 | 1 | 2 | 0 | 0 | 0 | 0 | 0 | 1 | 0 | 0 | 0 | 0 | 0 | 2 | 0 | 0 | 15 | 2 | 2 | 0 | 0 | 0 | 0 | 0 | 0 | 0 | 0 | 19 | 2 | 0 | 17 | 0 | 0 | 0 | 0 | 1 | 4 | 19 | 5 | 0 |
| G to A | 0 | 0 | 0 | 0 | 0 | 0 | 0 | 0 | 0 | 0 | 0 | 0 | 0 | 0 | 0 | 0 | 0 | 0 | 0 | 0 | 0 | 0 | 0 | 0 | 0 | 0 | 0 | 0 | 0 | 0 | 0 | 0 | 0 | 0 | 0 | 0 | 0 | 0 | 0 | 0 | 0 | 0 | 0 | 0 | 0 |
| Other | 0 | 0 | 0 | 0 | 0 | 0 | 0 | 0 | 0 | 0 | 0 | 1 | 0 | 0 | 0 | 0 | 0 | 0 | 0 | 0 | 0 | 0 | 0 | 1 | 0 | 0 | 0 | 0 | 0 | 1 | 0 | 0 | 0 | 0 | 0 | 0 | 0 | 0 | 0 | 0 | 0 | 0 | 0 | 0 | 0 |

| t | t | t | a | t | a | c | c | c | c | a | g | c | c | g | a | g | g | a | c | g | c | g | t | g | c | c | t | g | g | a | c | t | c | c | c | c | g | c | c | c | c | c | g | g |
|---|---|---|---|---|---|---|---|---|---|---|---|---|---|---|---|---|---|---|---|---|---|---|---|---|---|---|---|---|---|---|---|---|---|---|---|---|---|---|---|---|---|---|---|---|
| | | | | | | T | T | T | T | | | T | T | | | | | | T | | T | | | | T | T | | | | | | C | T | T | T | T | | | T | T | T | T | | |
| | | | | | | T | T | T | T | | | | T | | | | | | T | | | | | | | | | | | | | | | T | | | | | | | | T | | |
| | | | | | | T | T | T | T | | | | T | | | | | | T | | | | | | | | | | | | | | | T | | | | | | | | T | | |
| | | | | | | | | | | | | | | | | | | | | | | | | | | | | | | | T | | T | T | T | | | | T | T | T | | | |
| | | | | | | | | | | | | | | | | | | | | | | | | | | | | | | | T | | T | T | T | | | | T | T | T | | | |
| | | | | | | T | T | T | T | | | | T | | | | | | T | | | | | | | | | | | | | | | T | | | | | | | | T | | |
| | | | | | | T | T | T | T | | | | T | | | | | | T | | | | | | | | | | | | | | | T | | | | | | | | T | | |
| | | | | | | T | T | T | T | | | | T | | | | | | T | | | | | | | | | | | | | | | T | | | | | | | | T | | |
| | | | | | | T | T | T | T | | | | T | | | | | | T | | | | | | | | | | | | | | | T | | | | | | | | T | | |
| | | | | | | T | T | T | T | | | | T | | | | | | T | | | | | | | | | | | | | | | T | | | | | | | | T | | |
| | | | | | | T | T | T | T | | | | T | | | | | | T | | | | | | | | | | | | | | | T | | | | | | | | T | | |
| | | | | | | T | T | T | T | | | T | T | | | | | | | | | | | | T | | | | | | | | T | T | T | T | | | | | T | | | |
| | | | | | | T | T | T | T | | | | T | | | | | | T | | | | | | | | | | | | | | | T | | | | | | | | T | | |
| | | | | | | T | T | T | T | | | | T | | | | | | T | | | | | | | | | | | | | | | T | | | | | | | | T | | |
| | | | | | | T | T | T | T | | | | T | | | | | | T | | | | | | | | | | | | | | | T | | | | | | | | T | | |
| C | | | | | | | T | T | | | | | | | | | | | | | | | | | T | | | | | | T | | T | T | T | T | | T | T | T | T | T | | |
| | | | | | | T | T | T | T | | | | T | | | | | | T | | | | | | | | | | | | | | | T | | | | | | | | T | | |
| | | | | | | T | T | T | T | | | | T | | | | | | T | | | | | | | | | | | | | | | T | | | | | | | | T | | |
| | | | | | | T | T | T | T | | | | T | | | | | | T | | | | | | | | | | | | | | | T | | | | | | | | T | | |
| | | | | | | T | T | T | T | | | | T | | | | | | T | | | | | | | | | | | | | | | T | | | | | | | | T | | |
| 0 | 0 | 0 | 0 | 0 | 0 | 17 | 18 | 18 | 17 | 0 | 0 | 2 | 17 | 0 | 0 | 0 | 0 | 0 | 16 | 0 | 1 | 0 | 0 | 0 | 3 | 1 | 0 | 0 | 0 | 0 | 3 | 0 | 5 | 20 | 5 | 3 | 0 | 1 | 4 | 4 | 5 | 17 | 0 | 0 |
| 0 | 0 | 0 | 0 | 0 | 0 | 0 | 0 | 0 | 0 | 0 | 0 | 0 | 0 | 0 | 0 | 0 | 0 | 0 | 0 | 0 | 0 | 0 | 0 | 0 | 0 | 0 | 0 | 0 | 0 | 0 | 0 | 0 | 0 | 0 | 0 | 0 | 0 | 0 | 0 | 0 | 0 | 0 | 0 | 0 |
| 1 | 0 | 0 | 0 | 0 | 0 | 0 | 0 | 0 | 0 | 0 | 0 | 0 | 0 | 0 | 0 | 0 | 0 | 0 | 0 | 0 | 0 | 0 | 0 | 0 | 0 | 0 | 0 | 0 | 0 | 0 | 0 | 1 | 0 | 0 | 0 | 0 | 0 | 0 | 0 | 0 | 0 | 0 | 0 | 0 |

| a | g | a | c | c | c | c | c | a | a | a | c | c | t | t | c | c | c | a | c | a | c | c | a | c | a | c | c | a | c | c | c | g | g | c | g | a | t | g | c | c | g | a | g | c |
|---|---|---|---|---|---|---|---|---|---|---|---|---|---|---|---|---|---|---|---|---|---|---|---|---|---|---|---|---|---|---|---|---|---|---|---|---|---|---|---|---|---|---|---|---|
| | | | | T | T | T | T | | | | | T | | | T | T | T | | T | | | T | | | | | | | | | T | | | | | | | | | T | | | | |
| | | | | T | T | T | T | | | | | T | | | T | T | T | | T | | | T | | T | G | T | T | | | | | | | | | | | | | | | | | T |
| | | | | T | T | T | T | | | | | T | | | T | T | T | | T | | T | T | | T | | T | T | | | | | | | | | | | | | | | | | T |
| | | | T | T | T | T | T | | | | T | T | | | T | T | T | | T | | T | T | | T | | T | | | | T | | | | | | | | | | | | | | |
| | | | T | T | T | T | T | | | | T | T | | | T | T | T | | T | | T | T | | T | | T | | | | T | | | | | | | | | | | | | | |
| | | | | T | T | T | T | | | | | T | | | T | T | T | | T | | T | T | | T | | T | T | | | | | | | | | | | A | | | | | | T |
| | | | | T | T | T | T | | | | | T | | | T | T | T | | T | | T | T | | T | | T | T | | | | | | | | | G | | | | | | | | T |
| | | | | T | T | T | T | | | | | T | | | T | T | T | | T | | T | T | | T | | T | T | | | | | | | | | | | | | | | | | T |
| | | | | T | T | T | T | | | | | T | | | T | T | T | | T | | T | T | | T | | T | T | | | | | | | | | | | | | | | | | T |
| | | | | T | T | T | T | | | | | T | | | T | T | T | | T | | T | T | | T | | T | T | | | | | | | | | | | | | | | | | T |
| | | | | T | T | T | T | | | | | T | | | T | T | T | | T | | T | T | | T | | T | T | | | | | | | | | | | | | | | | | T |
| | | | T | T | T | T | T | | | | T | T | | | T | T | T | | | | | | | | | | | | | | | | | | | | | | | | | | | |
| | | | | T | T | T | T | | | | | T | | | T | T | T | | T | | T | T | | T | | T | T | | | | | | | | | | | | | | | | | T |
| | | | | T | T | T | T | | | | | T | | | T | T | T | | T | | T | T | | T | | T | T | | | | | | | | | | | | | | | | | T |
| | | | | T | T | T | T | | | | | T | | | T | T | T | | T | | T | T | | T | | T | T | | | | | | | | | | | | | | | | | T |
| | | | T | T | T | T | T | | | | T | T | | | T | T | T | | | | | T | | T | | T | T | | | T | | | | | | | | | | | | | | T |
| | | | | T | T | T | T | | | | | T | | | T | T | T | | T | | T | T | | T | | T | T | | | | | | | | | | | | | | | | | T |
| | | | | T | T | T | T | | | | | T | | | T | T | T | | T | | T | T | | T | | T | T | | | | | | | | | | | | | | | | | T |
| | | | | T | T | T | T | | | | | T | | | T | T | T | | T | | T | T | | T | | T | T | | | | | | | | | | | | | | | | | T |
| | | | | T | T | T | T | | | | | T | | | T | T | T | | T | | | T | | T | | T | T | | | | | | | | | | | | | | | | | T |
| 0 | 0 | 0 | 4 | 20 | 20 | 20 | 20 | 0 | 0 | 0 | 4 | 20 | 0 | 0 | 20 | 20 | 20 | 0 | 18 | 0 | 15 | 19 | 0 | 18 | 0 | 18 | 16 | 0 | 0 | 3 | 1 | 0 | 0 | 0 | 0 | 0 | 0 | 0 | 0 | 1 | 0 | 0 | 0 | 16 |
| 0 | 0 | 0 | 0 | 0 | 0 | 0 | 0 | 0 | 0 | 0 | 0 | 0 | 0 | 0 | 0 | 0 | 0 | 0 | 0 | 0 | 0 | 0 | 0 | 0 | 0 | 0 | 0 | 0 | 0 | 0 | 0 | 0 | 0 | 0 | 0 | 0 | 0 | 1 | 0 | 0 | 0 | 0 | 0 | 0 |
| 0 | 0 | 0 | 0 | 0 | 0 | 0 | 0 | 0 | 0 | 0 | 0 | 0 | 0 | 0 | 0 | 0 | 0 | 0 | 0 | 0 | 0 | 0 | 0 | 0 | 1 | 0 | 0 | 0 | 0 | 0 | 0 | 0 | 0 | 0 | 0 | 1 | 0 | 0 | 0 | 0 | 0 | 0 | 0 | 0 |

| g | c | c | t | g | t | g | t | c | a | t | c | t | g | c | a | g | g | a | g | a | t | c | c | t | g | g | c | c | c | a | g | a | t | g | t | a | c | g | g |
|---|---|---|---|---|---|---|---|---|---|---|---|---|---|---|---|---|---|---|---|---|---|---|---|---|---|---|---|---|---|---|---|---|---|---|---|---|---|---|---|
| C | | | | | | | | T | | | T | | | | | | | | | | | T | | | | | | T | T | | | | | | | | | | |
| | T | T | | | | | | T | | | T | | | | | | | | | | | T | | | | | | | | | | | | | | | T | | |
| | T | T | | | | | | T | | | T | | | | | | | | | | | T | | | | | | | | | | | | | | | T | | |
| | T | T | | | | | | T | | | T | | | T | | | | | | | | | | | | | | | | | | | | | | | | | |
| | T | T | | | | | | T | | | T | | | T | | | | | | | | | | | | | | | | | | | | | | | | | |
| | T | T | | | | | | T | | | T | | | | | | | | | | | | | | | | | | | | | | | | | | | | |
| | T | T | | | | | | T | | | T | | | | | | | | | | | T | | | | | | | | | | | | | | | T | | |
| | T | T | | | | | | T | | | T | | | | | | | | | | | T | | | | | | | | | | | | | | | T | | |
| | T | T | | | | | | T | | | T | | | | | | | | | | | T | | | | | | | | | | | | | | | T | | |
| | T | T | | | | | | T | | | T | | | | | | | | | | | T | | | | | | | | | | | | | | | T | | |
| | T | T | | | | | | T | | | T | | | | | | | | | | | T | | | | | | | | | | | | | | | T | | |
| A | T | T | | | | | | T | | | T | | | | | | | | | | | T | | | | | | | | | | | | | | | | | |
| | T | T | | | | | | T | | | T | | | | | | | | | | | T | | | | | | | | | | | | | | | T | | |
| | T | T | | | | | | T | | | T | | | | | | | | | | | T | | | | | | | | | | | | | | | T | | |
| | T | T | | | | | | T | | | T | | | T | | | | | | | | T | | | | | | | T | | | | | | | | T | | |
| | T | T | | | | | | T | | | T | | | | | | | | | | | T | | | | | | | | | | | | | | | T | | |
| | T | T | | | | | | T | | | T | | | | | | | | | | | | | | | | | | | | | | | | | | | | |
| | T | T | | | | | | T | | | T | | | | | | | | | | | T | | | | | | | | | | | | | | | T | | |
| | T | T | | | | | | T | | | T | | | | | | | | | | | T | | | | | | | | | | | | | | | T | | |
| 0 | # | # | 0 | 0 | 0 | 0 | 0 | 19 | 0 | 0 | 19 | 0 | 0 | 3 | 0 | 0 | 0 | 0 | 0 | 0 | 0 | 15 | 0 | 0 | 0 | 0 | 0 | 1 | 2 | 0 | 0 | 0 | 0 | 0 | 0 | 0 | 13 | 0 | 0 |
| 1 | 0 | 0 | 0 | 0 | 0 | 0 | 0 | 0 | 0 | 0 | 0 | 0 | 0 | 0 | 0 | 0 | 0 | 0 | 0 | 0 | 0 | 0 | 0 | 0 | 0 | 0 | 0 | 0 | 0 | 0 | 0 | 0 | 0 | 0 | 0 | 0 | 0 | 0 | 0 |
| 1 | 0 | 0 | 0 | 0 | 0 | 0 | 0 | 0 | 0 | 0 | 0 | 0 | 0 | 0 | 0 | 0 | 0 | 0 | 0 | 0 | 0 | 0 | 0 | 0 | 0 | 0 | 0 | 0 | 0 | 0 | 0 | 0 | 0 | 0 | 0 | 0 | 0 | 0 | 0 |

**93.8°C_vUNG-S302A + EGFP-P2A-stop**

| Clone number | Amino acid sequences of wild-type Us3 | | | | | | | | | | | | | | | | | | | | | | | | | | | | | | | | | | | | | | | | |
|---|---|---|---|---|---|---|---|---|---|---|---|---|---|---|---|---|---|---|---|---|---|---|---|---|---|---|---|---|---|---|---|---|---|---|---|---|---|---|---|---|---|
| | g | t | c | g | c | g | t | t | t | a | c | g | g | g | g | g | a | c | a | g | g | g | c | a | g | g | a | g | g | a | a | g | g | a | g | g | a | g | g | c | c | g |
| 1 | | | | | | | | | | | | | | | | | | | | | | | | | | | | | | | | | | | | | | | | | | |
| 2 | | | | | | | | | | | | | | | | | | | | | | | | | | | | | | | | | | | | | | | | | | |
| 3 | | | | | | | | | | | | | | | | | | | | | | | | | | | | | | | | | | | | | | | | | | |
| 4 | | | | | | | | | | | | | | | | | | | | | | | | | | | | | | | | | | | | | | | | | | |
| 5 | | | | | | | | | | | | | | | | | | | | | | | | | | | | | | | | | | | | | | | | | | |
| 6 | | | | | | | | | | | | | | | | | | | | | | | | | | | | | | | | | | | | | | | | | | |
| 7 | | | | | | | | | | | | | | | | | | | | | | | | | | | | | | | | | | | | | | | | | | |
| 8 | | | | | | | | | | | | | | | | | | | | | | | | | | | | | | | | | | | | | | | | | | |
| 9 | | | | | | | | | | | | | | | | | | | | | | | | | | | | | | | | | | | | | | | | | | |
| 10 | | | | | | | | | | | | | | | | | | | | | | | | | | | | | | | | | | | | | | | | | | |
| 11 | | | | | | | | | | | | | | | | | G | | | | | | | | | | | | | | | | | | | | | | | | | |
| 12 | | | | | | | | | | | | | | | | | | | | | | | | | | | | | | | | | | | | | | | | | | |
| 13 | | | | | | | | | | | | | | | | | | | | | | | | | | | | | | | | | | | | | | | | | | |
| 14 | | | | | T | | | | | | | | | | | | | | | | | | | | | | | | | | | | | | | | | | | | | |
| 15 | | | | | | | | | | | | | | | | | | | | | | | | | | | | | | | | | | | | | | | | | | |
| 16 | | | | | | | | | | | | | | | | | | | | | | | | | | | | | | | | | | | | | | | | | | |
| 17 | | | | | | | | | | | | | | | | | | | | | | | | | | | | | | | | | | | | | | | | | | |
| 18 | | | | | | | | | | | | | | | | | | | | | | | | | | | | | | | | | | | | | | | | | | |
| 19 | | | | | | | | | | | | | | | | | | | | | | | | | | | | | | | | | | | | | | | | | | |
| 20 | | | | | | | | | | | | | | | | | | | | | | | | | | | | | | | | | | | | | | | | | | |
| C to T | 0 | 0 | 0 | 0 | 1 | 0 | 0 | 0 | 0 | 0 | 0 | 0 | 0 | 0 | 0 | 0 | 0 | 0 | 0 | 0 | 0 | 0 | 0 | 0 | 0 | 0 | 0 | 0 | 0 | 0 | 0 | 0 | 0 | 0 | 0 | 0 | 0 | 0 | 0 | 0 | 0 | 0 |
| G to A | 0 | 0 | 0 | 0 | 0 | 0 | 0 | 0 | 0 | 0 | 0 | 0 | 0 | 0 | 0 | 0 | 0 | 0 | 0 | 0 | 0 | 0 | 0 | 0 | 0 | 0 | 0 | 0 | 0 | 0 | 0 | 0 | 0 | 0 | 0 | 0 | 0 | 0 | 0 | 0 | 0 | 0 |
| Other | 0 | 0 | 0 | 0 | 0 | 0 | 0 | 0 | 0 | 0 | 0 | 0 | 0 | 0 | 0 | 0 | 1 | 0 | 0 | 0 | 0 | 0 | 0 | 0 | 0 | 0 | 0 | 0 | 0 | 0 | 0 | 0 | 0 | 0 | 0 | 0 | 0 | 0 | 0 | 0 | 0 | 0 |

| t | c | c | c | g | c | c | g | g | a | g | a | c | a | a | a | g | c | c | g | t | c | c | c | g | g | g | t | g | t | t | t | c | c | t | c | a | t | g | g | c | c | c | c | t |
|---|---|---|---|---|---|---|---|---|---|---|---|---|---|---|---|---|---|---|---|---|---|---|---|---|---|---|---|---|---|---|---|---|---|---|---|---|---|---|---|---|---|---|---|---|
| 0 | 0 | 0 | 0 | 0 | 0 | 0 | 0 | 0 | 0 | 0 | 0 | 0 | 0 | 0 | 0 | 0 | 0 | 0 | 0 | 0 | 0 | 0 | 0 | 0 | 0 | 0 | 0 | 0 | 0 | 0 | 0 | 0 | 0 | 0 | 0 | 0 | 0 | 0 | 0 | 0 | 0 | 0 | 0 | 0 |
| 0 | 0 | 0 | 0 | 0 | 0 | 0 | 0 | 0 | 0 | 0 | 0 | 0 | 0 | 0 | 0 | 0 | 0 | 0 | 0 | 0 | 0 | 0 | 0 | 0 | 0 | 0 | 0 | 0 | 0 | 0 | 0 | 0 | 0 | 0 | 0 | 0 | 0 | 0 | 0 | 0 | 0 | 0 | 0 | 0 |
| 0 | 0 | 0 | 0 | 0 | 0 | 0 | 0 | 0 | 0 | 0 | 0 | 0 | 0 | 0 | 0 | 0 | 0 | 0 | 0 | 0 | 0 | 0 | 0 | 0 | 0 | 0 | 0 | 0 | 0 | 0 | 0 | 0 | 0 | 0 | 0 | 0 | 0 | 0 | 0 | 0 | 0 | 0 | 0 | 0 |

| t | t | t | a | t | a | c | c | c | c | a | g | c | c | g | a | g | g | a | c | g | c | g | t | g | c | c | t | g | g | a | c | t | c | c | c | c | g | c | c | c | c | c | g | g |
|---|---|---|---|---|---|---|---|---|---|---|---|---|---|---|---|---|---|---|---|---|---|---|---|---|---|---|---|---|---|---|---|---|---|---|---|---|---|---|---|---|---|---|---|---|
| 0 | 0 | 0 | 0 | 0 | 0 | 0 | 0 | 0 | 0 | 0 | 0 | 0 | 0 | 0 | 0 | 0 | 0 | 0 | 0 | 0 | 0 | 0 | 0 | 0 | 0 | 0 | 0 | 0 | 0 | 0 | 0 | 0 | 0 | 0 | 0 | 0 | 0 | 0 | 0 | 0 | 0 | 0 | 0 | 0 |
| 0 | 0 | 0 | 0 | 0 | 0 | 0 | 0 | 0 | 0 | 0 | 0 | 0 | 0 | 0 | 0 | 0 | 0 | 0 | 0 | 0 | 0 | 0 | 0 | 0 | 0 | 0 | 0 | 0 | 0 | 0 | 0 | 0 | 0 | 0 | 0 | 0 | 0 | 0 | 0 | 0 | 0 | 0 | 0 | 0 |
| 0 | 0 | 0 | 0 | 0 | 0 | 0 | 0 | 0 | 0 | 0 | 0 | 0 | 0 | 0 | 0 | 0 | 0 | 0 | 0 | 0 | 0 | 0 | 0 | 0 | 0 | 0 | 0 | 0 | 0 | 0 | 0 | 0 | 0 | 0 | 0 | 0 | 0 | 0 | 0 | 0 | 0 | 0 | 0 | 0 |

| a | g | a | c | c | c | c | c | a | a | a | c | c | t | t | c | c | c | a | c | a | c | c | a | c | a | c | c | a | c | c | c | g | g | c | g | a | t | g | c | c | g | a | g | c |
|---|---|---|---|---|---|---|---|---|---|---|---|---|---|---|---|---|---|---|---|---|---|---|---|---|---|---|---|---|---|---|---|---|---|---|---|---|---|---|---|---|---|---|---|---|
| 0 | 0 | 0 | 0 | 0 | 0 | 0 | 0 | 0 | 0 | 0 | 0 | 0 | 0 | 0 | 0 | 0 | 0 | 0 | 0 | 0 | 0 | 0 | 0 | 0 | 0 | 0 | 0 | 0 | 0 | 0 | 0 | 0 | 0 | 0 | 0 | 0 | 0 | 0 | 0 | 0 | 0 | 0 | 0 | 0 |
| 0 | 0 | 0 | 0 | 0 | 0 | 0 | 0 | 0 | 0 | 0 | 0 | 0 | 0 | 0 | 0 | 0 | 0 | 0 | 0 | 0 | 0 | 0 | 0 | 0 | 0 | 0 | 0 | 0 | 0 | 0 | 0 | 0 | 0 | 0 | 0 | 0 | 0 | 0 | 0 | 0 | 0 | 0 | 0 | 0 |
| 0 | 0 | 0 | 0 | 0 | 0 | 0 | 0 | 0 | 0 | 0 | 0 | 0 | 0 | 0 | 0 | 0 | 0 | 0 | 0 | 0 | 0 | 0 | 0 | 0 | 0 | 0 | 0 | 0 | 0 | 0 | 0 | 0 | 0 | 0 | 0 | 0 | 0 | 0 | 0 | 0 | 0 | 0 | 0 | 0 |

| g | c | c | t | g | t | g | t | c | a | t | c | t | g | c | a | g | g | a | g | a | t | c | c | t | g | g | c | c | c | a | g | a | t | g | t | a | c | g | g |
|---|---|---|---|---|---|---|---|---|---|---|---|---|---|---|---|---|---|---|---|---|---|---|---|---|---|---|---|---|---|---|---|---|---|---|---|---|---|---|---|
| 0 | 0 | 0 | 0 | 0 | 0 | 0 | 0 | 0 | 0 | 0 | 0 | 0 | 0 | 0 | 0 | 0 | 0 | 0 | 0 | 0 | 0 | 0 | 0 | 0 | 0 | 0 | 0 | 0 | 0 | 0 | 0 | 0 | 0 | 0 | 0 | 0 | 0 | 0 | 0 |
| 0 | 0 | 0 | 0 | 0 | 0 | 0 | 0 | 0 | 0 | 0 | 0 | 0 | 0 | 0 | 0 | 0 | 0 | 0 | 0 | 0 | 0 | 0 | 0 | 0 | 0 | 0 | 0 | 0 | 0 | 0 | 0 | 0 | 0 | 0 | 0 | 0 | 0 | 0 | 0 |
| 0 | 0 | 0 | 0 | 0 | 0 | 0 | 0 | 0 | 0 | 0 | 0 | 0 | 0 | 0 | 0 | 0 | 0 | 0 | 0 | 0 | 0 | 0 | 0 | 0 | 0 | 0 | 0 | 0 | 0 | 0 | 0 | 0 | 0 | 0 | 0 | 0 | 0 | 0 | 0 |

**93.8°C_vUNG-S302A + EGFP-P2A-hAPOBEC1**

| Clone number | Amino acid sequences of wild-type Us3 | | | | | | | | | | | | | | | | | | | | | | | | | | | | | | | | | | | | | | | | |
|---|---|---|---|---|---|---|---|---|---|---|---|---|---|---|---|---|---|---|---|---|---|---|---|---|---|---|---|---|---|---|---|---|---|---|---|---|---|---|---|---|---|
| | g | t | c | g | c | g | t | t | t | a | c | g | g | g | g | g | a | c | a | g | g | g | c | a | g | g | a | g | g | a | a | g | g | a | g | g | a | g | g | c | c | g |
| 1 | | | | | | | | | | | | | | | | | | | | | | | | | | | | | | | | | | | | | | | | | | |
| 2 | | | | | | | | | | | | | | | | | | | | | | | | | | | | | | | | | | | | | | | | | | |
| 3 | | | | | | | | | | | | | | | | | | | | | | | | | | | | | | | | | | | | | | | | | | |
| 4 | | | | | | | | | | | | | | | | | | | | | | | | | | | | | | | | | | | | | | | | | | |
| 5 | | | | | | | | | | | | | | | | | | | | | | | | | | | | | | | | | | | | | | | | | | |
| 6 | | | | | | | | | | | | | | | | | | | | | | | | | | | | | | | | | | | | | | | | | | |
| 7 | | | | | | | | | | | | | | | | | | | | | | | | | | | | | | | | | | | | | | | | | | |
| 8 | | | | | | | | | | | | | | | | | | | | | | | | | | | | | | | | | | | | | | | | | | |
| 9 | | | | | | | | | | | | | | | | | | | | | | | | | | | | | | | | | | | | | | | | | | |
| 10 | | | | | | | | | | | | | | | | | | | | | | | | | | | | | | | | | | | | | | | | | | |
| 11 | | | | | | | | | | | | | | | | | | | | | | | | | | | | | | | | | | | | | | | | | | |
| 12 | | | | | | | | | | | | | | | | | | | | | | | | | | | | | | | | | | | | | | | | | | |
| 13 | | | | | | | | | | | | | | | | | | | | | | | | | | | | | | | | | | | | | | | | | | |
| 14 | | | | | | | | | | | | | | | | | | | | | | | | | | | | | | | | | | | | | | | | | | |
| 15 | | | | | | | | | | | | | | | | | | | | | | | | | | | | | | | | | | | | | | | | | | |
| 16 | | | | | | | | | | | | | | | | | | | | | | | | | | | | | | | | | | | | | | | | | | |
| 17 | | | | | | | | | | | | | | | | | | | | | | | | | | | | | | | | | | | | | | | | | | |
| 18 | | | | | | | | | | | | | | | | | | | | | | | | | | | | | | | | | | | | | | | | | | |
| 19 | | | | | | | | | | | | | | | | | | | | | | | | | | | | | | | | | | | | | | | | | | |
| 20 | | | A | | | | | | | | | | | | | | | | | | | | | | | | | | | | | | | | | | | | | | | |
| C to T | 0 | 0 | 0 | 0 | 0 | 0 | 0 | 0 | 0 | 0 | 0 | 0 | 0 | 0 | 0 | 0 | 0 | 0 | 0 | 0 | 0 | 0 | 0 | 0 | 0 | 0 | 0 | 0 | 0 | 0 | 0 | 0 | 0 | 0 | 0 | 0 | 0 | 0 | 0 | 0 | 0 | 0 |
| G to A | 0 | 0 | 0 | 0 | 0 | 0 | 0 | 0 | 0 | 0 | 0 | 0 | 0 | 0 | 0 | 0 | 0 | 0 | 0 | 0 | 0 | 0 | 0 | 0 | 0 | 0 | 0 | 0 | 0 | 0 | 0 | 0 | 0 | 0 | 0 | 0 | 0 | 0 | 0 | 0 | 0 | 0 |
| Other | 0 | 0 | 1 | 0 | 0 | 0 | 0 | 0 | 0 | 0 | 0 | 0 | 0 | 0 | 0 | 0 | 0 | 0 | 0 | 0 | 0 | 0 | 0 | 0 | 0 | 0 | 0 | 0 | 0 | 0 | 0 | 0 | 0 | 0 | 0 | 0 | 0 | 0 | 0 | 0 | 0 | 0 |

| Clone number | t | c | c | c | g | c | c | g | g | a | g | a | c | a | a | a | g | c | c | g | t | c | c | c | g | g | g | t | g | t | t | t | c | c | t | c | a | t | g | g | c | c | c | c | t |
|---|---|---|---|---|---|---|---|---|---|---|---|---|---|---|---|---|---|---|---|---|---|---|---|---|---|---|---|---|---|---|---|---|---|---|---|---|---|---|---|---|---|---|---|---|---|
| 1 | | | | | | T | | | | | | | | | | | | | | | | | | | | | | | | | | | | | | | | | | | | | | | |
| 2 | | | | | | | | | | | | | | | | | | | | | | | | | | | | | | | | | | | | | | | | | | | | | |
| 3 | | | | | | | | | | | | | | | | | | | | | | | | | | | | | | | | | | | | | | | | | | | | | |
| 4 | | | | | | | | | | | | | | | | | | | | | | | | | | | | | | | | | | | | | | | | | | | | | |
| 5 | | | | | | | | | | | | | | | | | | | | | | | | | | | | | | | | | | | | | | | | | | | | | |
| 6 | | | | | | | | | | | | | | | | | | | | | | | | | | | | | | | | | | | | | | | | | | | | | |
| 7 | | | | | | | | | | | | | | | | | | | | | | | | | | | | | | | | | | | | | | | | | | | | | |
| 8 | | | | | | | | | | | | | | | | | | | | | | | | | | | | | | | | | | | | | | | | | | | | | |
| 9 | | | | | | | | | | | | | | | | | | | | | | | | | | | | | | | | | | | | | | | | | | | | | |
| 10 | | | | | | | | | | | | | | | | | | | | | | | | | | | | | | | | | | | | | | | | | | | | | |
| 11 | | | | | | | | | | | | G | | | | | | | | | | | | | | | | | | | | | | | | | | | | | | | | | |
| 12 | | | | | | | | | | | | | | | | | | | | | | | | | | | | | | | | | | | | | | | | | | | | | |
| 13 | | | | | | | | | | | | | | | | | | | | | | | | | | | | | | | | | | | | | | | | | | | | | |
| 14 | | | | | | | | | | | | | | | | | | | | | | | | | | | | | | | | | | | | | | | | | | | | | |
| 15 | | | | | | | | | | | | | | | | | | | | | | | | | | | | | | | | | | | | | | | | | | | | | |
| 16 | | | | | | | | | | | | | | | | | | | | | | | | | | | | | | | | | | | | | | | | | | | | | |
| 17 | | | | | | | | | | | | | | | | | | | | | | | | | | | | | | | | | | | | | | | | | | | | | |
| 18 | | | | | | | | | | | | | | | | | | | | | | | | | | | | | | | | | | | | | | | | | | | | | |
| 19 | | | | | | | | | | | | | | | | | | | | | | | | | | | | | | | | | | | | | | | | | | | | | |
| 20 | | | | | | | | | | | | | | | | | | | | | | | | | | | | | | | | | | | | | | | A | | | | | | |
| C to T | 0 | 0 | 0 | 0 | 0 | 1 | 0 | 0 | 0 | 0 | 0 | 0 | 0 | 0 | 0 | 0 | 0 | 0 | 0 | 0 | 0 | 0 | 0 | 0 | 0 | 0 | 0 | 0 | 0 | 0 | 0 | 0 | 0 | 0 | 0 | 0 | 0 | 0 | 0 | 0 | 0 | 0 | 0 | 0 | 0 |
| G to A | 0 | 0 | 0 | 0 | 0 | 0 | 0 | 0 | 0 | 0 | 0 | 0 | 0 | 0 | 0 | 0 | 0 | 0 | 0 | 0 | 0 | 0 | 0 | 0 | 0 | 0 | 0 | 0 | 0 | 0 | 0 | 0 | 0 | 0 | 0 | 0 | 0 | 0 | 1 | 0 | 0 | 0 | 0 | 0 | 0 |
| Other | 0 | 0 | 0 | 0 | 0 | 0 | 0 | 0 | 0 | 0 | 0 | 1 | 0 | 0 | 0 | 0 | 0 | 0 | 0 | 0 | 0 | 0 | 0 | 0 | 0 | 0 | 0 | 0 | 0 | 0 | 0 | 0 | 0 | 0 | 0 | 0 | 0 | 0 | 0 | 0 | 0 | 0 | 0 | 0 | 0 |

| | | | | | | | | | | | | | | | | | | | | | | | | | | | | | | | | | | | | | | | | | | | | |
|---|---|---|---|---|---|---|---|---|---|---|---|---|---|---|---|---|---|---|---|---|---|---|---|---|---|---|---|---|---|---|---|---|---|---|---|---|---|---|---|---|---|---|---|---|
| t | t | t | a | t | a | c | c | c | c | a | g | c | c | g | a | g | g | a | c | g | c | g | t | g | c | c | t | g | g | a | c | t | c | c | c | c | g | c | c | c | c | c | g | g |
| | | | | | | | | | | | | | | | | | | | | | | | | | | | | | | | | | | | | | | | | | T | | | |
| 0 | 0 | 0 | 0 | 0 | 0 | 0 | 0 | 0 | 0 | 0 | 0 | 0 | 0 | 0 | 0 | 0 | 0 | 0 | 0 | 0 | 0 | 0 | 0 | 0 | 0 | 0 | 0 | 0 | 0 | 0 | 0 | 0 | 0 | 0 | 0 | 0 | 0 | 0 | 0 | 0 | 1 | 0 | 0 | 0 |
| 0 | 0 | 0 | 0 | 0 | 0 | 0 | 0 | 0 | 0 | 0 | 0 | 0 | 0 | 0 | 0 | 0 | 0 | 0 | 0 | 0 | 0 | 0 | 0 | 0 | 0 | 0 | 0 | 0 | 0 | 0 | 0 | 0 | 0 | 0 | 0 | 0 | 0 | 0 | 0 | 0 | 0 | 0 | 0 | 0 |
| 0 | 0 | 0 | 0 | 0 | 0 | 0 | 0 | 0 | 0 | 0 | 0 | 0 | 0 | 0 | 0 | 0 | 0 | 0 | 0 | 0 | 0 | 0 | 0 | 0 | 0 | 0 | 0 | 0 | 0 | 0 | 0 | 0 | 0 | 0 | 0 | 0 | 0 | 0 | 0 | 0 | 0 | 0 | 0 | 0 |

| | | | | | | | | | | | | | | | | | | | | | | | | | | | | | | | | | | | | | | | | | | | | |
|---|---|---|---|---|---|---|---|---|---|---|---|---|---|---|---|---|---|---|---|---|---|---|---|---|---|---|---|---|---|---|---|---|---|---|---|---|---|---|---|---|---|---|---|---|
| a | g | a | c | c | c | c | c | a | a | a | c | c | t | t | c | c | c | a | c | a | c | c | a | c | a | c | c | a | c | c | c | g | g | c | g | a | t | g | c | c | g | a | g | c |
| | | | | | | | | | | | | | | | | | | | | | | | | | | | | | | | | | | T | | | | | | | | | | |
| | | | | | | | | | | | | | | | | | | | | | | | | | | | | | | | | | | | | | | | | T | | | | |
| | | | | | | | T | | | | | | | | | | | | | | | | | | | | | | | | | | | | | | | | | | | | | |
| 0 | 0 | 0 | 0 | 0 | 0 | 0 | 1 | 0 | 0 | 0 | 0 | 0 | 0 | 0 | 0 | 0 | 0 | 0 | 0 | 0 | 0 | 0 | 0 | 0 | 0 | 0 | 0 | 0 | 0 | 0 | 0 | 0 | 0 | 1 | 0 | 0 | 0 | 0 | 0 | 1 | 0 | 0 | 0 | 0 |
| 0 | 0 | 0 | 0 | 0 | 0 | 0 | 0 | 0 | 0 | 0 | 0 | 0 | 0 | 0 | 0 | 0 | 0 | 0 | 0 | 0 | 0 | 0 | 0 | 0 | 0 | 0 | 0 | 0 | 0 | 0 | 0 | 0 | 0 | 0 | 0 | 0 | 0 | 0 | 0 | 0 | 0 | 0 | 0 | 0 |
| 0 | 0 | 0 | 0 | 0 | 0 | 0 | 0 | 0 | 0 | 0 | 0 | 0 | 0 | 0 | 0 | 0 | 0 | 0 | 0 | 0 | 0 | 0 | 0 | 0 | 0 | 0 | 0 | 0 | 0 | 0 | 0 | 0 | 0 | 0 | 0 | 0 | 0 | 0 | 0 | 0 | 0 | 0 | 0 | 0 |

| g | c | c | t | g | t | g | t | c | a | t | c | t | g | c | a | g | g | a | g | a | t | c | c | t | g | g | c | c | c | a | g | a | t | g | t | a | c | g | g |
|---|---|---|---|---|---|---|---|---|---|---|---|---|---|---|---|---|---|---|---|---|---|---|---|---|---|---|---|---|---|---|---|---|---|---|---|---|---|---|---|
| | | | C | | | | | | | | | | | | | | | | | | | | | | | | | | | | | | | | | | | | |
| 0 | 0 | 0 | 0 | 0 | 0 | 0 | 0 | 0 | 0 | 0 | 0 | 0 | 0 | 0 | 0 | 0 | 0 | 0 | 0 | 0 | 0 | 0 | 0 | 0 | 0 | 0 | 0 | 0 | 0 | 0 | 0 | 0 | 0 | 0 | 0 | 0 | 0 | 0 | 0 |
| 0 | 0 | 0 | 0 | 0 | 0 | 0 | 0 | 0 | 0 | 0 | 0 | 0 | 0 | 0 | 0 | 0 | 0 | 0 | 0 | 0 | 0 | 0 | 0 | 0 | 0 | 0 | 0 | 0 | 0 | 0 | 0 | 0 | 0 | 0 | 0 | 0 | 0 | 0 | 0 |
| 0 | 0 | 0 | 1 | 0 | 0 | 0 | 0 | 0 | 0 | 0 | 0 | 0 | 0 | 0 | 0 | 0 | 0 | 0 | 0 | 0 | 0 | 0 | 0 | 0 | 0 | 0 | 0 | 0 | 0 | 0 | 0 | 0 | 0 | 0 | 0 | 0 | 0 | 0 | 0 |

**93.8°C_vUNG-SA-repair + EGFP-P2A-stop**

| Clone number | Amino acid sequences of wild-type Us3 | | | | | | | | | | | | | | | | | | | | | | | | | | | | | | | | | | | | | | | | |
|---|---|---|---|---|---|---|---|---|---|---|---|---|---|---|---|---|---|---|---|---|---|---|---|---|---|---|---|---|---|---|---|---|---|---|---|---|---|---|---|---|---|
| | g | t | c | g | c | g | t | t | t | a | c | g | g | g | g | g | a | c | a | g | g | g | c | a | g | g | a | g | g | a | a | g | g | a | g | g | a | g | g | c | c | g |
| 1 | | | | | | | | | | | | | | | | | | | | | | | | | | | | | | | | | | | | | | | | | | |
| 2 | | | | | | | | | | | | | | | | | | | | | | | | | | | | | | | | | | | | | | | | | | |
| 3 | | | | | | | | C | | | | | | | | | | | | | | | | | | | | | | | | | | | | | | | | | | |
| 4 | | | | | | | | | | | | | | | | | | | | | | | | | | | | | | | | | | | | | | | | | | |
| 5 | | | | | | | | | | | | | | | | | | | | | | | | | | | | | | | | | | | | | | | | | | |
| 6 | | | | | | | | | | | | | | | | | | | | | | | | | | | | | | | | | | | | | | | | | | |
| 7 | | | | | | | | | | | | | | | | | | | | | | | | | | | | | | | | | | | | | | | | | | |
| 8 | | | | | | | | | | | | | | | | | | | | | | | | | | | | | | | | | | | | | | | | | | |
| 9 | | | | | | | | | | | | | | | | | | | | | | | | | | | | | | | | | | | | | | | | | | |
| 10 | | | | | | | | | | | | | | | | | | | | | | | | | | | | | | | | | | | | | | | | | | |
| 11 | | | | | | | | | | | | | | | | | | | | | | | | | | | | | | | | | | | | | | | | | | |
| 12 | | | | | | | | | | | | | | | | | | | | | | | | | | | | | | | | | | | | | | | | | | |
| 13 | | | | | | | | | | | | | | | | | | | | | | | | | | | | | | | | | | | | | | | | | | |
| 14 | | | | | | | | | | | | | | | | | | | | | | | | | | | | | | | | | | | | | | | | | | |
| 15 | | | | | | | | | | | | | | | | | | | | | | | | | | | | | | | | | | | | | | | | | | |
| 16 | | | | | | | | | | | | | | | | | | | | | | | | | | | | | | | | | | | | | | | | | | |
| 17 | | | | | | | | | | | | | | | | | | | | | | | | | | | | | | | | | | | | | | | | | | |
| 18 | | | | | | | | | | | | | | | | | | | | | | | | | | | | | | | | | | | | | | | | | | |
| 19 | | | | | | | | | | | | | | | | | | | | | | | | | | | | | | | | | | | | | | | | | | |
| 20 | | | | | | | | | | | | | | | | | | | | | | | | | | | | | | | | | | | | | | | | | | |
| C to T | 0 | 0 | 0 | 0 | 0 | 0 | 0 | 0 | 0 | 0 | 0 | 0 | 0 | 0 | 0 | 0 | 0 | 0 | 0 | 0 | 0 | 0 | 0 | 0 | 0 | 0 | 0 | 0 | 0 | 0 | 0 | 0 | 0 | 0 | 0 | 0 | 0 | 0 | 0 | 0 | 0 | 0 |
| G to A | 0 | 0 | 0 | 0 | 0 | 0 | 0 | 0 | 0 | 0 | 0 | 0 | 0 | 0 | 0 | 0 | 0 | 0 | 0 | 0 | 0 | 0 | 0 | 0 | 0 | 0 | 0 | 0 | 0 | 0 | 0 | 0 | 0 | 0 | 0 | 0 | 0 | 0 | 0 | 0 | 0 | 0 |
| Other | 0 | 0 | 0 | 0 | 0 | 0 | 0 | 1 | 0 | 0 | 0 | 0 | 0 | 0 | 0 | 0 | 0 | 0 | 0 | 0 | 0 | 0 | 0 | 0 | 0 | 0 | 0 | 0 | 0 | 0 | 0 | 0 | 0 | 0 | 0 | 0 | 0 | 0 | 0 | 0 | 0 | 0 |

t c c c g c c g g a g a c a a a g c c g t c c c g g g t g t t t c c t c a t g g c c c c t

T

A

A

T

G

0 0 0 0 0 0 0 0 0 0 0 0 0 0 0 0 0 0 0 0 0 0 0 0 0 0 0 0 0 0 0 0 0 0 0 0 0 0 0 0 0 0 0 1 0

0 0 0 0 0 0 0 0 0 0 0 0 0 0 0 0 0 0 0 0 0 0 0 0 0 0 0 0 0 0 0 0 0 0 0 0 0 0 0 1 0 0 0 0 0

0 0 0 0 0 0 0 1 0 0 0 0 0 0 1 0 0 0 0 0 0 0 0 0 0 0 0 0 0 1 0 0 0 0 0 0 0 0 0 0 0 0 0 0 0

t t t a t a c c c c a g c c g a g g a c g c g t g c c t g g a c t c c c c g c c c c c g g

T G

A

0 0 0 0 0 0 0 0 0 0 0 0 0 1 0 0 0 0 0 0 0 0 0 0 0 0 0 0 0 0 0 0 0 0 0 0 0 0 0 0 0 0 0 0 0

0 0 0 0 0 0 0 0 0 0 0 0 0 0 0 0 0 0 0 0 0 0 0 0 1 0 0 0 0 0 0 0 0 0 0 0 0 0 0 0 0 0 0 0 0

0 0 0 0 0 0 0 0 0 0 0 0 0 0 0 0 0 0 0 0 0 0 0 0 0 0 0 0 0 0 1 0 0 0 0 0 0 0 0 0 0 0 0 0 0

| | | | | | | | | | | | | | | | | | | | | | | | | | | | | | | | | | | | | | | | | | | | | |
|---|---|---|---|---|---|---|---|---|---|---|---|---|---|---|---|---|---|---|---|---|---|---|---|---|---|---|---|---|---|---|---|---|---|---|---|---|---|---|---|---|---|---|---|---|
| a | g | a | c | c | c | c | c | a | a | a | c | c | t | t | c | c | c | a | c | a | c | c | a | c | a | c | c | a | c | c | c | g | g | c | g | a | t | g | c | c | g | a | g | c |
| | | | | | | | | | | | | | | | | | | | | | | | | | | | | | | | | | | T | | | | | | | | | | |
| 0 | 0 | 0 | 0 | 0 | 0 | 0 | 0 | 0 | 0 | 0 | 0 | 0 | 0 | 0 | 0 | 0 | 0 | 0 | 0 | 0 | 0 | 0 | 0 | 0 | 0 | 0 | 0 | 0 | 0 | 0 | 0 | 0 | 0 | 1 | 0 | 0 | 0 | 0 | 0 | 0 | 0 | 0 | 0 | 0 |
| 0 | 0 | 0 | 0 | 0 | 0 | 0 | 0 | 0 | 0 | 0 | 0 | 0 | 0 | 0 | 0 | 0 | 0 | 0 | 0 | 0 | 0 | 0 | 0 | 0 | 0 | 0 | 0 | 0 | 0 | 0 | 0 | 0 | 0 | 0 | 0 | 0 | 0 | 0 | 0 | 0 | 0 | 0 | 0 | 0 |
| 0 | 0 | 0 | 0 | 0 | 0 | 0 | 0 | 0 | 0 | 0 | 0 | 0 | 0 | 0 | 0 | 0 | 0 | 0 | 0 | 0 | 0 | 0 | 0 | 0 | 0 | 0 | 0 | 0 | 0 | 0 | 0 | 0 | 0 | 0 | 0 | 0 | 0 | 0 | 0 | 0 | 0 | 0 | 0 | 0 |

| | | | | | | | | | | | | | | | | | | | | | | | | | | | | | | | | | | | | | | | |
|---|---|---|---|---|---|---|---|---|---|---|---|---|---|---|---|---|---|---|---|---|---|---|---|---|---|---|---|---|---|---|---|---|---|---|---|---|---|---|---|
| g | c | c | t | g | t | g | t | c | a | t | c | t | g | c | a | g | g | a | g | a | t | c | c | t | g | g | c | c | c | a | g | a | t | g | t | a | c | g | g |
| 0 | 0 | 0 | 0 | 0 | 0 | 0 | 0 | 0 | 0 | 0 | 0 | 0 | 0 | 0 | 0 | 0 | 0 | 0 | 0 | 0 | 0 | 0 | 0 | 0 | 0 | 0 | 0 | 0 | 0 | 0 | 0 | 0 | 0 | 0 | 0 | 0 | 0 | 0 | 0 |
| 0 | 0 | 0 | 0 | 0 | 0 | 0 | 0 | 0 | 0 | 0 | 0 | 0 | 0 | 0 | 0 | 0 | 0 | 0 | 0 | 0 | 0 | 0 | 0 | 0 | 0 | 0 | 0 | 0 | 0 | 0 | 0 | 0 | 0 | 0 | 0 | 0 | 0 | 0 | 0 |
| 0 | 0 | 0 | 0 | 0 | 0 | 0 | 0 | 0 | 0 | 0 | 0 | 0 | 0 | 0 | 0 | 0 | 0 | 0 | 0 | 0 | 0 | 0 | 0 | 0 | 0 | 0 | 0 | 0 | 0 | 0 | 0 | 0 | 0 | 0 | 0 | 0 | 0 | 0 | 0 |

**Supplementary Table 3. Oligonucleotide sequences and DNA template for the construction of plasmids.**

| Constructed plasmid | Oligonucleotide sequence (5′-3′) | PCR DNA template | Recipient plasmid |
|---|---|---|---|
| pcDNA-EGFP-P2A-stop | 5′-GCGGATCCGCCACCATGGTGAGCAAGGGCGAGGA-3′ | pEGFP-C1 (Clontech) | pcDNA3.1(+) (Thermo Fisher Scientific) |
| | 5′-GCGAATTCAGGCCCGGGGTTTTCTTCAACATCTCCTGCTTGCTTTAACAGAGAGAAGTTCGTGGCTCCGCTTCCCTTGTACAGCTCGTCCATGC-3′ | | |
| pcDNA-EGFP-P2A-hAPOBEC1 | 5′-CCCCGGGCCTGAATTCATGACTTCTGAGAAAGGTCC-3′ | p3flag-hA1 (This study) | pcDNA-EGFP-P2A (This study) |
| | 5′-CCGCCACTGTGCTGGATATCTCATCTCCAAGCCACAGAAG-3′ | | |
| pEGFP-mA1 | 5′-GAATTCACCATGGCTCAGAAGGAAGAGGC-3′ | pGEM-mA1 (This study) | pEGFP-C2 (Clontech) |
| | 5′-TCTCGAGTCATTTCAACCCTGTAGCCCAAA-3′ | | |
| pRetroX-TRE3G-hAPOBEC1-3xflag | 5′-GCGGATCCGCCACCATGACTTCTGAGAAAGGTCC-3′ | pcDNA-hAPOBEC1 (This study) | pRetroX-TRE3G (TaKaRa) |
| | 5′-GCGAATTCTCACTTGTCATCGTCATCCTTGTAATCGATGTCATGATCTTTATAATCACCGTCATGGTCTTTGTAGTCTCTCCAAGCCACAGAAGGAT-3′ | | |
| pRetroX-TRE3G-mAPOBEC1-3xflag | 5′-GCGGATCCGCCACCATGAGTTCCGAGACAGGCCC-3′ | pEGFP-mA1 (This study) | pRetroX-TRE3G (TaKaRa) |
| | 5′-GCGAATTCTCACTTGTCATCGTCATCCTTGTAATCGATGTCA TGATCTTTATAATCACCGTCATGGTCTTTGTAGTCTTTCAACCCTGTAGCCC-3′ | | |
| pBS-UGI-flag | 5′-ATGTCGCGGCCGCGCCACCATGACAAATTTATCTGACAT -3′ | UGI-pFERp[45] | pBluescript II KS(+) (Stratagene) |
| | 5′GCGAATTCTTACTTGTCATCGTCGTCCTTGTAATCTAACATTTTAATTTTATTTTC-3′ | | |
| pAAV-UGI | 5′-GCGAATTCGCCACCATGACAAATTTATCTGACAT-3′ | pBS-UGI-flag (This study) | pAAV-ZsGreen1 (TaKaRa) |
| | 5′-GCGGATCCTTATAACATTTTAATTTTAT-3′ | | |

**Supplementary Table 4. Oligonucleotide sequences for the construction of recombinant viruses.**

| Recombinant virus | Oligonucleotide sequence (5′-3′) | Plasmid DNA template | *E. coli* GS1873 containing HSV-1 BAC |
|---|---|---|---|
| vUNG-S53A | 5'-CGGCGCATCTAATGATGCCGCGACGGAAACCCGTCCGGGTGCGGGGGGCGAACCGGCCGCAGGATGACGACGATAAGTAGGG-3' | pEP-KanS[48] | *E. coli* GS1783 /pYEbac102Cre[27, 46] |
| | 5'-CCGCCGGCCCTGACGAGCGACAGGCGGCCGGTTCGCCCCCCGCACCCGGACGGGTTTCCGCAACCAATTAACCAATTCTGATTAG-3' | | |
| vUNG-S302A | 5'-GGACCCTCGGGTCCATTGCGTCCTCAAGTTTTCGCACCCGGCTCCCCTCTCCAAGGTTCCGTTAGGATGACGACGATAAGTAGGG-3' | pEP-KanS[48] | *E. coli* GS1783 /pYEbac102Cre[27, 46] |
| | 5'-GGAAATGCTGGCATGTTCCGAACGGAACCTTGGAGAGGGGAGCCGGGTGCGAAAACTTGAGGACAACCAATTAACCAATTCTGATTAG-3' | | |
| vUNG-Q177L/D178N | 5'-TATTGCACCCCCGACGAGGTGCGCGTGGTTATCATCGGCCTGAACCCATATCACCACCCCGGCAGGATGACGACGATAAGTAGGG-3' | pEP-KanS[48] | *E. coli* GS1783 /pYEbac102Cre[27, 46] |
| | 5'-AACGCAAGTCCGTGCGCCTGGCCGGGGTGGTGATATGGGTTCAGGCCGATGATAACCACGCGCCAACCAATTAACCAATTCTGATTAG-3' | | |
| Flag-vUNG-S53A /FlagvUNG-S302A /Flag-vUNG-Q177L/D178N | 5'-TCTCGAGACCCGGTCGATTTCACCCATCGACTGGTCGGTTGACTACAAAGACGATGACGACAAGTGAAAGGCAGGATGACGACGATAAGTAGGG-3' | pEP-KanS[48] | *E. coli* GS1783 containing the vUNG -S53A, -S302A or -Q177L/D178N genome (This study) |
| | 5'-AGCCCCCACAGACGAAAACCCCGGACGTCGATGCCTTTCACTTGTCGTCATCGTCTTTGTAGTCAACCGACCCAACCAATTAACCAATTCTGATTAG-3' | | |

| | | | |
|---|---|---|---|
| vUNG-SA-repair /Flag-vUNG-SA-repair | 5’-GGACCCTCGGGTCCATTGCGTCCTCAAGTTTTCGCACCCGGACCCCCTCT CCAAGGTTCCGTTAGGATGACGACGATAAGTAGGG-3’ | pEP-KanS[48] | *E. coli* GS1783 containing the vUNG-S302A or Flag-vUNG-S302A genome (This study) |
| | 5’-GGAAATGCTGGCATGTTCCGAACGGAACCTTGGAGAGGGGGTCCGGGTG CGAAAACTTGAGGACAACCAATTAACCAATTCTGATTAG-3’ | | |
| | 5’-GGACCCTCGGGTCCATTGCGTCCTCAAGTTTTCGCACCCGTCGCCCCTCT CCAAGGTTCCGTTAGGATGACGACGATAAGTAGGG-3’ | | |
| | 5’-GGAAATGCTGGCATGTTCCGAACGGAACCTTGGAGAGGGGCGACGGGTG CGAAAACTTGAGGACAACCAATTAACCAATTCTGATTAG-3’ | | |
| vUNG-QL/DN-repair /Flag-vUNG-QL/DN-repair | 5’-TATTGCACCCCCGACGAGGTGCGCGTGGTTATCATCGGCCAGGACCCATA TCACCACCCCGGAGGATGACGACGATAAGTAGGG-3’ | pEP-KanS[48] | *E. coli* GS1783 containing the vUNG-Q177L/D178N or Flag-vUNG-Q177L/D178N genome (This study) |
| | 5’-AACGCAAGTCCGTGCGCCTGGCCGGGGTGGTGATATGGGTCCTGGCCGA TGATAACCACGCCAACCAATTAACCAATTCTGATTAG-3’ | | |
